# PD-1 remodels SHP2 dynamics and drives the non-catalytic inhibition of T cell activation

**DOI:** 10.64898/2026.05.20.725130

**Authors:** Panyu Fei, Peng Jiao, Jie Gao, Hui Chen, Yong Zhang, Lu Chen, Peng Wu, Lina Chen, Salvatore Valvo, Michael L. Dustin, Jizhong Lou, Chun Zhou, Wei Chen

## Abstract

The inhibitory receptor programmed cell death protein 1 (PD-1) suppresses T cell activation primarily by recruiting the src homology 2 domain-containing phosphatase 2 (SHP2), thereby unleashing its phosphatase activity. However, a purely enzymatic model struggles to fully explain the rapid and robust disruption of proximal T cell receptor (TCR) and CD28 signaling. Here, by integrating X-ray crystallography, single-molecule magnetic tweezers, supported lipid bilayers, TIRF imaging, liquid–liquid phase separation, and cellular assays, we uncover a dual-layered inhibitory mechanism of the PD-1/SHP2 axis. Mechanistically, we determine the crystal structure of SHP2 in complex with dually phosphorylated PD-1 cytoplasmic motifs (pPD-1), revealing a ligand-induced open conformation, which is distinct from the oncogenic E76K-associated state. We further directly resolve the closed-to-open conformational dynamics of individual SHP2 molecules, characterized by an 8-nm transition amplitude, and demonstrate that pPD-1 accelerates the transition by at least 94-fold. Crucially, the pPD-1-mediated highly open conformation exposes the tandem SH2 (t-SH2) domains to function as a biophysical barrier. Independent of phosphatase activity, these domains directly dismantle pTCR and pCD28 signaling condensates to suppress T cell activation. Together, our findings establish a model in which pPD-1 remodels SHP2 structural dynamics and drives a non-catalytic inhibitory mechanism of TCR and CD28 signaling, suggesting next-generation immunotherapy design to reverse T cell suppression.

## Introduction

First cloned and characterized in 1992 (human)^1,2^ and in 1993 (mouse),^3^ SHP2 is a non-receptor protein tyrosine phosphatase encoded by the *PTPN11* gene. It contains two SH2 (N-SH2 and C-SH2) domains, a protein tyrosine phosphatase (PTP) catalytic domain, and a C-terminal tail. In its basal state, SHP2 adopts a closed, autoinhibited conformation wherein the N-SH2 domain occludes the PTP catalytic site.^4–6^

PD-1 was first cloned and identified in 1992,^7^ containing an extracellular IgV-like domain, a transmembrane domain, and a cytoplasmic tail. PD-1 has been validated as a central immune checkpoint receptor expressed on T cells and is significantly activated in tumor microenvironments and during chronic stimulation upon the interaction with its ligands, PD-L1^8^ or PD-L2.^9^ Consequently, PD-1 plays a critical role in restraining T cell antitumor responses and has emerged as a major clinical target for the treatment of cancer and autoimmune diseases.^10–13^ In the context of immune regulation, SHP2 is a central mediator of PD-1-driven inhibitory signaling in T cells. For decades, the prevailing dogma has postulated that PD-1 suppresses T cell activation almost exclusively by recruiting SHP2 to enzymatically dephosphorylate proximal signaling molecules, such as the CD28 co-stimulatory receptor and the TCR–CD3 complex.^14–16^ Mechanistically, upon ligand engagement, the immunoreceptor tyrosine-based inhibitory motif (ITIM) and immunoreceptor tyrosine-based switch motif (ITSM) within the PD-1 cytoplasmic tail become phosphorylated (pITIM and pITSM), enabling the recruitment and activation of SHP2, unleashing its phosphatase activity.^15,17^

Previous structural studies have shown that, in the absence of PD-1 engagement, SHP2 adopts an autoinhibited conformation in the cytosol, in which the N-SH2 domain occludes the PTP catalytic site, rendering the phosphatase inactive.^4,18^ Gain-of-function SHP2 mutations located at the N-SH2–PTP interface stabilize an open conformation of SHP2,^19^ which is markedly distinct from the autoinhibited structure. Moreover, structural analyses of the N-SH2 and C-SH2 domains in complex with phosphorylated pITIM and pITSM motifs^20^ reveal that the N-SH2 domain preferentially engages pITIM, whereas the C-SH2 domain binds pITSM. Together, these findings support a prevailing model in which SHP2 undergoes a conformational transition from a closed, autoinhibited state to an open, active state upon binding to phosphorylated PD-1, thereby exposing the PTP catalytic domain and permitting access to tyrosine-phosphorylated substrates. This conformational transition is thought to be essential for SHP2 activation and the consequent attenuation of T cell activation.

Although the closed-to-open conformational transition model of SHP2 implies a central role for conformational dynamics in SHP2 function, prior studies have primarily captured static structures of SHP2 alone or in truncated SHP2–PD-1 complexes.^4,20^ As a result, the real-time conformational dynamics of full-length SHP2 following PD-1 phosphorylation, and their contribution to T cell inhibitory signaling, remain largely unexplored.

Earlier investigations revealed that SHP2 is rapidly recruited to PD-1 within the first 10–12 seconds following TCR and PD-1 stimulation, whereas statistically significant dephosphorylation of downstream TCR signaling molecules^15^ and attenuation of Jurkat T cell Ca^2+^ signaling^21^ are not observed until approximately 2 minutes later. Despite the well-established recognition of SHP2’s catalytic function in inhibiting T cells, this purely enzymatic model struggles to explain the immediate spatial disruption of immune synapses. This temporal discrepancy prompts speculation regarding the existence of a non-catalytic mode of SHP2-mediated T cell activation inhibition.

Recent studies have indicated that SHP2 can regulate signaling cascades through biomolecular condensates formed via liquid–liquid phase separation (LLPS). For instance, phosphorylated receptor tyrosine kinases (RTKs), such as FGFR2, undergo LLPS with SHP2 through phosphorylation-dependent multivalent SH2–pY interactions, thereby forming membraneless compartments that fine-tune functional outputs.^22^ Collectively, these lines of evidence lead us to explore whether SHP2 exerts non-catalytic inhibition via molecular multivalent interactions at the T cell immune synapse, beyond its canonical catalytic role.

To resolve this mechanism, it is necessary to bridge the gap between single-molecule conformational dynamics and mesoscale cellular assemblies. In this study, we combine multidimensional methodologies to delineate how PD-1 phosphorylation dynamically reshapes the conformational dynamics and energy landscape of SHP2, transforming it into a non-catalytic condensate disruptor to rapidly dissolve pTCR and pCD28 signaling condensates, thereby providing a comprehensive understanding of immune checkpoint signaling.

## Results

### Crystal structures of SHP2–pPD-1 complex reveal a pronounced open conformation of SHP2

Although structural analyses of the N-SH2 and C-SH2 domains in complex with phosphorylated pITIM and pITSM motifs have been reported previously,^20^ to elucidate how PD-1 remodels the conformation of SHP2, we crystallized core SHP2 (residues: 1–525; N-SH2, C-SH2, and PTP) in complex with phosphorylated PD-1 ITIM and ITSM motifs (Figures 1A–1D and S1, and Table S1).

**Figure 1.**
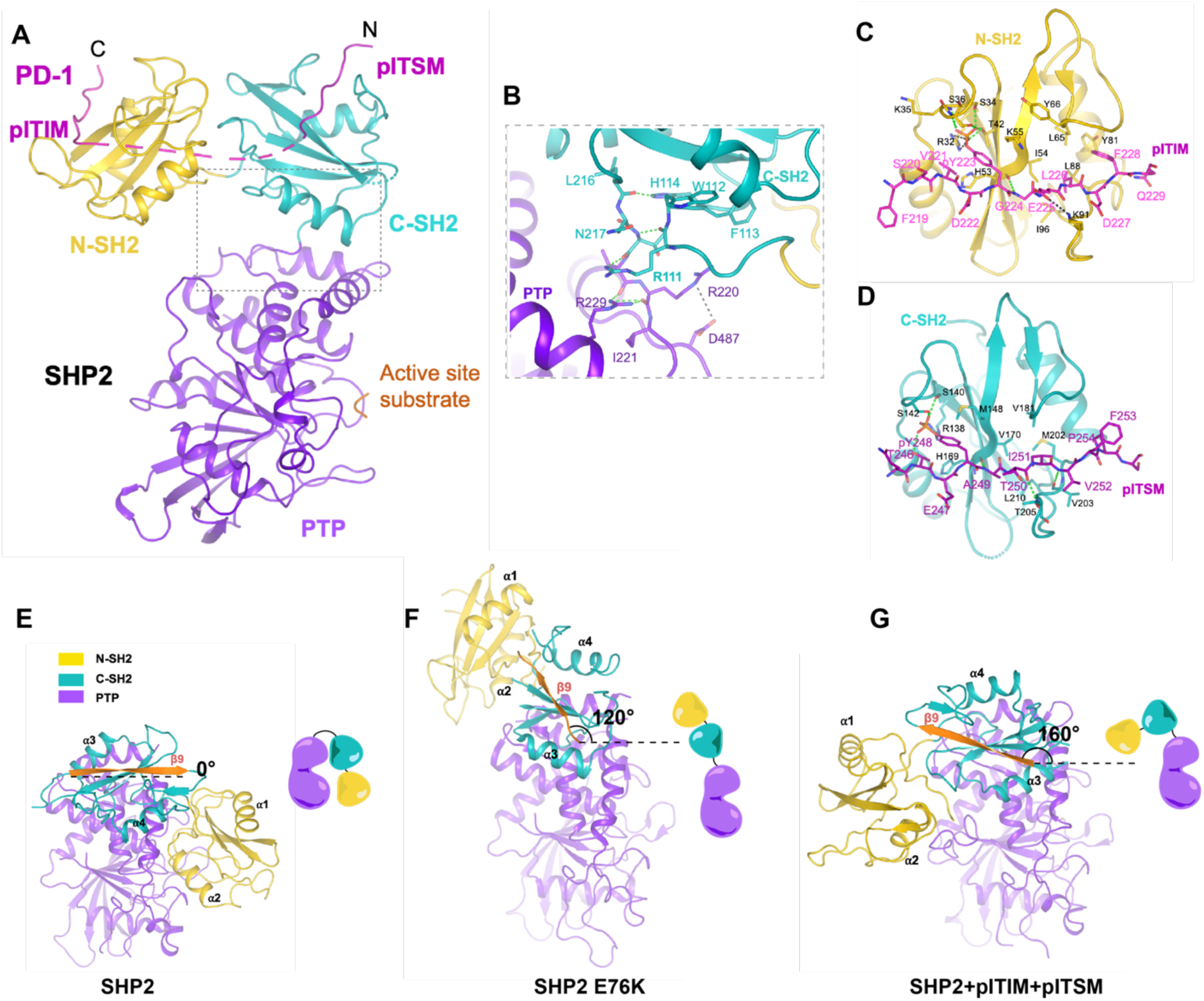
Crystal structures of SHP2 in complex with phosphorylated PD-1 motifs reveal a prominent open conformation of SHP2. (A) Ribbon representation of SHP2–PD1 complex (PDB: 21BJ, in this study). N-SH2, C-SH2, and PTP are colored yellow, teal, and purple-blue, respectively. PD-1 motifs bound to SH2 domains are colored in magenta. A three-residue fragment of the phosphorylated PD1 motifs (built as ApYA) is also observed in the PTP catalytic site. (B) Residues at the C-SH2–PTP domain interface are displayed in the dashed rectangle in (A), green dashed lines indicate hydrogen bonds, and grey dashed lines indicate salt bridges. The view is flipped around 180 degrees to show the detailed interactions. (C and D) Diagram showing detained interactions between N-SH2–pITIM (C) and C-SH2–pITSM (D), green dashed lines indicate hydrogen bonds, grey dashed lines indicate salt bridges. (E–G) Crystal structures of SHP2 (PDB: 2SHP) (D), or SHP2 E76K (PDB: 6CRF) (E), or SHP2 bound to phosphorylated PD-1 motifs (PDB: 21BJ, from this study) (F) show that SHP2 undergoes significant conformational changes. Each structure is accompanied by a cartoon diagram of SHP2 on the right to illustrate its conformational changes. The three structures were aligned based on the PTP domain; β9 in the C-SH2 domain is colored orange to highlight the orientation of the C-SH2 domain, and the rotation degree is labelled. SHP2 in (A–D) and (G) is core SHP2 (residues: 1–525) carrying the C459S mutation, and SHP2 in (E) and (F) is core SHP2 (residues: 1–525). pITIM motif (residues: 218–234, sequence: VFSVDpYGELDFQWREKT), and pITSM motif (residues: 242–256, sequence: VPEQTEpYATIVFPSG). See also Figures 1 and S2.

The 2.6 Å crystal structure reveals that SHP2 adopts a markedly open conformation, with the PTP domain fully exposed and the pITIM and pITSM motifs bound to the N-SH2 and C-SH2 domains, respectively (Figure 1A). Notably, the linker region between the C-SH2 and PTP domains adopts an extended conformation. This conformation is stabilized by a localized interaction network involving R229 from the PTP domain, together with R111 and H114 from the flexible linker connecting the N-SH2 and C-SH2 domains. Specifically, R229 forms two hydrogen bonds with the main-chain carbonyl groups of T219 and R220. R111 stacks against R229 and forms two hydrogen bonds with the main-chain atoms of T218. This network is further anchored by H114, which hydrogen-bonds to the backbone carbonyl of L216 (Figure 1B).

The two phosphorylated PD-1 motifs bind in nearly parallel orientations but in opposite directions (Figures 1C and 1D). In the N-SH2 domain, the phosphate group of pY223 forms salt bridges with R32 and is further stabilized by hydrogen bonds with S34, S36, and T42. The aromatic ring of pY223 is sandwiched between PD-1 V211 and the alkyl chain of SHP2 K55. The amide group of G224 forms a main-chain hydrogen bond with the carbonyl group of SHP2 H53. In addition, the negatively charged side chain of pITIM E225 is complemented by K91, while two hydrophobic residues, L226 and F228, pack against a hydrophobic surface on N-SH2 formed by I54, L65, Y81, and L88 (Figure 1C).

In the C-SH2 domain, the phosphate group of pY248 is coordinated by SHP2 R138 and S142 and further stabilized by interactions with S140 and pITSM T246. The aromatic ring of pY248 packs against the side chain of SHP2 M148. PD-1 residues A249, T250, and V252 form main-chain hydrogen bonds with SHP2 H169, T205, and V203, respectively. Additionally, the hydrophobic residues I251 and V252 of pITSM engage a hydrophobic patch in the C-SH2 domain formed by V170, V181, M202, L210, and V203 (Figure 1D). Overall, the interactions between PD-1 motifs and the N-SH2 and C-SH2 domains of SHP2 are highly analogous to those observed in the previously reported N-SH2–pITIM crystal structure and C-SH2–pITSM NMR structure.^20^

According to previous reports, apo SHP2 adopts a closed conformation in which the N-SH2 domain occludes the catalytic site of the PTP domain (Figure 1E),^4^ whereas the disease-associated E76K mutant adopts an open conformation (Figure 1F).^19^ Structural comparison of these SHP2 conformations with the SHP2–pPD-1 motif complex (Figures 1E–1G) demonstrates that phosphorylated PD-1 not only promotes SHP2 opening but also induces more extensive domain rearrangements. In particular, the C-SH2 domain undergoes an approximately 160° rotation, repositioning the N-SH2 domain to the opposite side of the PTP catalytic site (Figure 1G). In contrast, the rotation of the C-SH2 domain in the E76K mutant is around 100° (Figure 1F).

In addition, structural analysis unexpectedly revealed a dimeric SHP2–pPD-1 motif complex in which the N-SH2 domain of one SHP2 molecule forms a stable interaction with the PTP domain of another (Figures 2A and S2B). The presence of this dimeric architecture may further stabilize the pPD-1-driven open confirmation of SHP2.

### Single-molecule magnetic tweezers capture the dynamic conformational changes of individual SHP2 molecules

The pronounced conformational transition of SHP2 revealed by our structures raised fundamental questions regarding how the closed-to-open transition occurs and whether the dynamics can be directly captured and quantified. To address these questions, we first performed steered molecular dynamics (SMD) simulations.^23^ Upon application of force, the earliest and most prominent conformational change observed in SHP2 was the dissociation of the N-SH2 domain from the PTP domain, with an extension amplitude of approximately 8 nm (Figure 2A).

**Figure 2.**
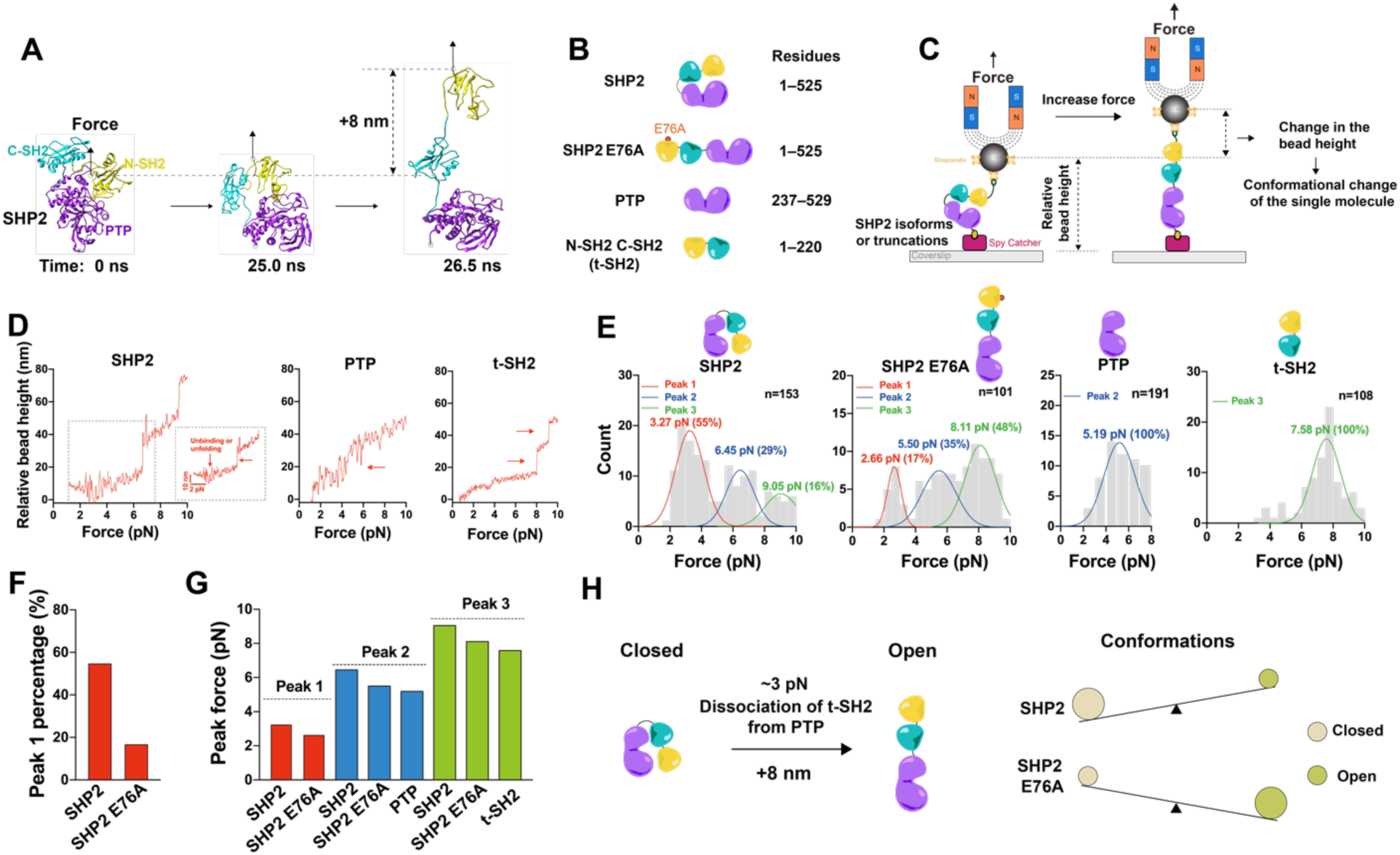
Individual SHP2 molecules undergo a conformational opening with a magnitude of 8 nm. (A) Using steered molecular dynamics (SMD) assay to reveal the conformational change of SHP2 under force. (B) The constructed SHP2 isoforms shown in the single-molecule magnetic tweezers (MT) experiments. (C) The diagram of the MT experiments to capture the conformational change of SHP2 at the single-molecule level. (D) Example force-relative bead height curves of the MT experiments for different SHP2 isoforms, unbinding or unfolding events with the molecule were indicated by red arrows (*N* ≥ 5, *n* is labelled). (E) The unbinding or unfolding forces histograms for different SHP2 isoforms, distinct force peaks were shown by the Gaussian fit, and the number of single molecules revealed was indicated. (F) Comparison of the first force peak percentage between SHP2 and SHP2 E76A. (G) Comparison of the peak forces in the three peaks, respectively, for different SHP2 isoforms. (H) Diagram of the conformational switch between the closed and open states of SHP2, where SHP2 tends to be closed, whereas SHP2 E76A tends to be open, and the applied ∼3 pN force promotes the switch from the closed to the open state in SHP2. See also Figure S3–S5.

Motivated by these results, we next sought to experimentally validate the conformational dynamics. To this end, we utilized the single-molecule magnetic tweezers (MT) previously established in our laboratory^23–25^ to capture conformational changes in SHP2 at single-molecule resolution. In MT experiments, conformational changes of individual proteins can arise either from interdomain dissociation or from unfolding within individual domains, which are demonstrated by the bead height. To distinguish between these possibilities, we constructed SHP2, the SHP2 E76A mutant, the truncations of the isolated PTP domain, or N-SH2 C-SH2 (t-SH2) domains (Figure 2B) for MT measurements (Figure 2C). By comparing SHP2 with the mutant or truncations, we were able to pinpoint the structural origin of each observed conformational change.

Upon application of force, all constructs exhibited single or multiple conformational transitions attributable to either interdomain dissociation or domain unfolding (Figure 2D). Next, we focused on the low-force regime (0–10 pN) to identify the earliest conformational transition in SHP2. Both SHP2 and the SHP2 E76A mutant displayed three distinct force peaks at approximately 3, 5, and 8 pN, corresponding to three independent conformational changes within individual molecules. Combining these data with the SHP2 crystal structure (Figure 1D) and SMD simulations, we attributed the first change occurring at ∼3 pN to the dissociation of the N-SH2 domain from the PTP domain.

Notably, compared with SHP2, the fraction of molecules exhibiting this first change was markedly reduced in the SHP2 E76A mutant, from 55% to 17%, which is in agreement with E76A favoring an open conformation. This observation further supports the assignment of the ∼3 pN transition to N-SH2–PTP domain dissociation (Figures 2E and 2F). In addition, quantitative analyses revealed that the second (∼5 pN) and third (∼8 pN) changes arise from the unfolding of the PTP domain and the t-SH2 domains, respectively (Figures 2D–2G).

Together, these results demonstrate that SHP2 and the SHP2 E76A mutant predominantly adopt closed and open conformations, respectively, and that a modest force of ∼3 pN is sufficient to drive the closed-to-open conformational transition in SHP2 by disrupting the N-SH2–PTP interaction (Figure 2H).

### Individual SHP2 molecules display reversible conformational switching

Given that SHP2 might undergo reversible switching between open and closed conformations to precisely tune PD-1-mediated inhibitory signaling in T cells, and that a low-level force of ∼3 pN is sufficient to drive the closed-to-open conformational transition (Figure 2), we therefore asked whether the dynamic conformational switching of individual SHP2 molecules could be directly captured and quantified. To this end, we held individual SHP2 molecules under a constant force of 3.5 pN in MT experiments to monitor the bead height trajectories over time, which represent the conformational dynamics of single SHP2 molecules. We then included the specific allosteric inhibitor SHP099^26^ as a critical negative control, which acts as a molecular glue to stabilize interactions among the N-SH2, C-SH2, and PTP domains (Figure S3), and tracked the conformational behaviors of the same SHP2 molecule before and after the introduction of SHP099 (Figure 3A).

**Figure 3.**
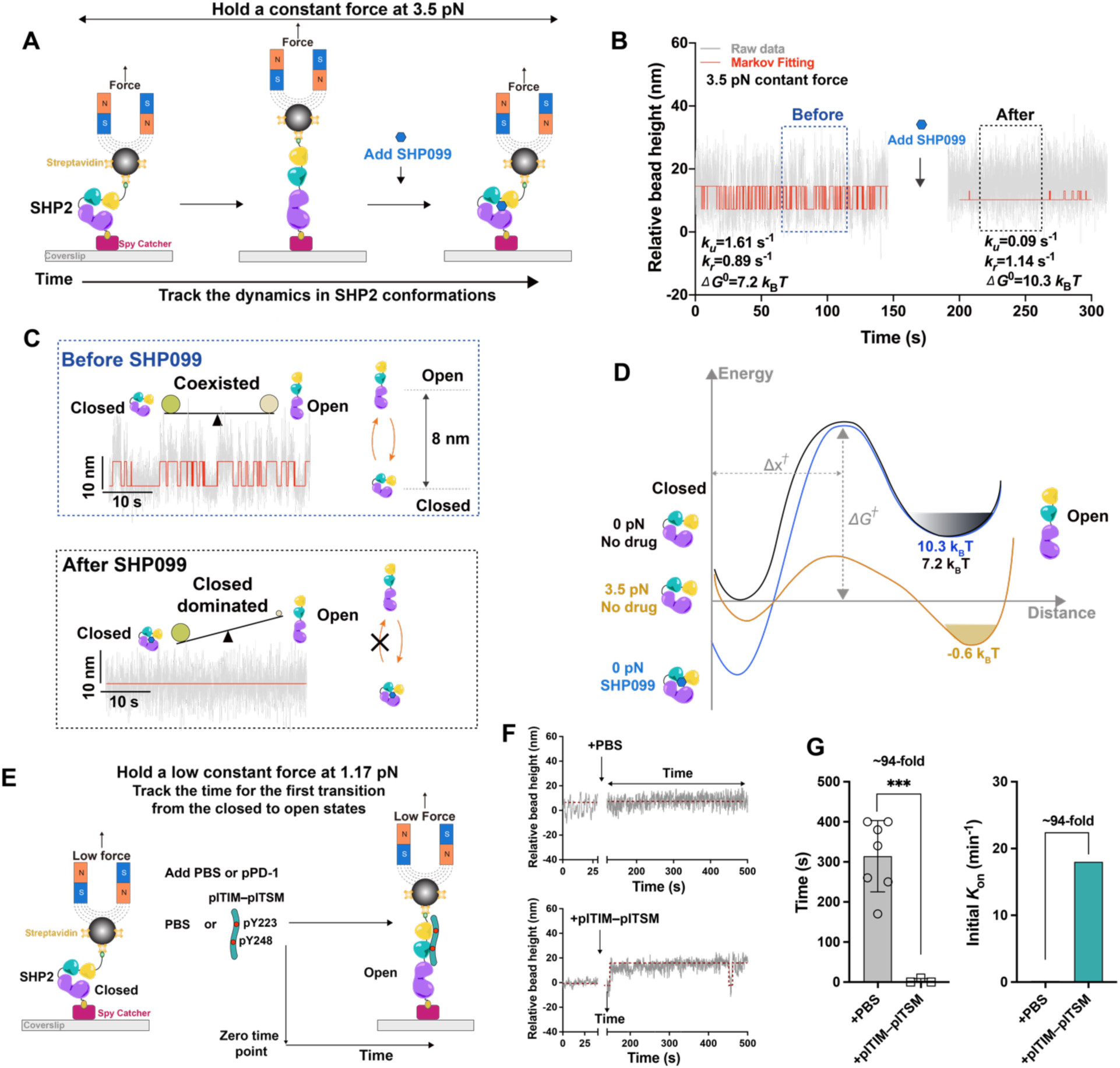
The dynamic conformational switch from closed to open states of individual SHP2 molecules is accelerated by phosphorylated PD-1. (A and B) The diagram (A) and results (B) of using MT to capture the dynamic conformational switch of individual SHP2 molecules under constant force with or without the presence of SHP099. (C) The zoom-in data in (B) as indicated by the dashed rectangle. (D) The energy landscapes of SHP2 with or without SHP099 or force. (E–G) The diagram (E), representative data (F), and results (G) of using MT to track the time and *K*_on_ for the first transition from closed to open state of SHP2 with PBS or pPD-1. The time tracking starts from around 100 s and stops at 500 s (*N* ≥ 3, *n* = 7 for PBS, *n* = 3 for pPD-1). The *K*_on_ is the reciprocal of the average time. SHP2 shown here is core SHP2 (residues: 1–525). Data in (G) were presented as mean ± SD. The significance of the difference was determined by an unpaired two-tailed Student’s *t*-test: \*\*\**P* ≤ 0.001. See also Figures 3–S5.

Before the addition of SHP099, we successfully captured that SHP2 exhibited frequent transitions between two states separated by an extension change of ∼8 nm (Figures 3B and 3C), consistent with the conformational difference predicted by SMD simulations (Figure 2A). Kinetic analysis yielded transition rates of 1.61 s^-1^ for opening and 0.89 s^-1^ for closing, corresponding to a free energy difference of 7.2 *k*_B_*T* required to promote SHP2 from the closed to the open state (Figures 3B and 3D).

Upon addition of SHP099 to the same molecule, conformational switching was strongly suppressed and occurred only rarely, with a markedly reduced extension amplitude (Figures 3B and 3C). Quantitatively, the opening rate decreased to 0.09 s^-1^, while the closing rate increased to 1.14 s^-1^. The free energy required to drive the closed-to-open transition increased substantially to 10.3 *k*_B_*T* (Figures 3B and 3D). Based on these measurements, we constructed the energy landscapes^27^ governing SHP2 conformational switching (Figure 3D). Without an inhibitor, although 7.2 *k*_B_*T* is required to promote SHP2 opening under zero force, the application of 3.5 pN effectively lowers the free energy difference to -0.6 *k*_B_*T*, enabling spontaneous transitions and rendering the energy barrier readily surmountable. In contrast, SHP099 stabilizes the closed conformation by increasing the free energy for opening to 10.3 *k*_B_*T*, thereby strongly disfavoring the open state.

To further validate the inhibitory effect of SHP099 at the single-molecule level, we sequentially added SHP099, washed it out, and finally introduced DMSO as a vehicle control during MT measurements while continuously tracking SHP2 conformational dynamics (Figure S4A). Prior to SHP099 treatment, SHP2 exhibited frequent switching between closed and open states. Following SHP099 addition, switching was abolished, with the closed conformation becoming dominant. Notably, after removal of SHP099, conformational switching was restored, indicating that the inhibitory effect was reversible. Finally, the addition of DMSO alone did not induce significant changes in SHP2 conformational behavior, confirming that the observed allosteric inhibition arose specifically from SHP099 (Figure S4B).

We previously found that the t-SH2 domains undergo two distinct conformational changes, arising from the sequential unfolding of the two SH2 domains under an applied force of ∼8 pN (Figure 2E). Here, we sought to directly capture the dynamic unfolding and refolding transitions of these domains. To this end, we held the force constant at 8.0 pN, 8.5 pN, or 9.0 pN in MT experiments to monitor the conformational dynamics of individual SHP2 molecules (Figure S5A). At 8.0 pN, we observed three distinct conformational states, designated states 1, 2, and 3. State 1 corresponds to both N-SH2 and C-SH2 domains remaining folded. State 2 represents the unfolding of a single SH2 domain (assigned here to N-SH2) while the other SH2 domain (assigned to C-SH2) remains folded. State 3 corresponds to the unfolding of both SH2 domains. Kinetic analysis revealed transitions among these states (1↔2 and 2↔3), with state 1 being the dominant population at 8.0 pN (Figure S5B, top). When the force was increased by 0.5 pN to 8.5 pN, we observed a pronounced increase in transitions from state 1 to state 2, resulting in state 2 becoming the dominant conformation (Figure S5B, middle). Further increasing the force to 9.0 pN led to a marked enhancement in the population of state 3, indicating extensive unfolding of both SH2 domains (Figure S5B, bottom).

### Phosphorylated PD-1 significantly accelerates the closed-to-open switch of individual SHP2 molecules

After characterizing the conformational dynamics of SHP2, we next asked how its main upstream signaling receptor, PD-1, regulates these dynamics, particularly the transition from the closed to the open state. To address this question, we tethered a single SHP2 molecule in the MT and applied a constant force of 1.17 pN, a regime that minimally perturbs the energy barrier for the closed-to-open transition. Under such low-force conditions, SHP2 alone would be expected to remain predominantly closed and to open only rarely over extended timescales.

We then introduced PBS or PD-1-derived phosphorylated motifs, pITIM–pITSM, into the system and measured the time to the first closed-to-open transition of the single SHP2 molecule (Figure 3E). In the presence of pITIM–pITSM, SHP2 transitioned from the closed to the open state within ∼10 s. By contrast, the opening time increased by approximately 94-fold with PBS. These measurements defined distinct opening rates, demonstrating striking differences in the ability of the phosphorylated PD-1 to promote the closed-to-open conformational switch of SHP2 in an extremely short time (Figure 3F).

### PD-1 phosphorylated states control the affinity of SHP2 and PD-1

The observation that SHP2 opening is accelerated by phosphorylated PD-1 suggests substantial differences in the binding kinetics between SHP2 and PD-1. Because molecular affinity is determined by both association and dissociation rates, we hypothesized that the affinity between SHP2 and PD-1 varies with PD-1’s phosphorylation states. Based on our structural analyses and prior studies,^15,20^ we identified two key regulatory tyrosines of Y223 in the ITIM motif and Y248 in the ITSM motif.

To test this hypothesis, we constructed and purified the full-length PD-1 intracellular domain (ICD) as well as two tyrosine mutants, PD-1 Y223F and PD-1 Y248F. These proteins were incubated in vitro with either full-length Lck or the truncated kinase domain of Lck in the presence of ATP. Phosphorylation at Y223 and/or Y248 was validated by high-resolution mass spectrometry (Figure S6A). Using this in vitro reconstitution system, we found that the Lck kinase domain, but not full-length Lck, efficiently phosphorylated all PD-1 isoforms (Figure S6B). This approach yielded four PD-1 species: fully phosphorylated PD-1 (pPD-1), singly phosphorylated PD-1 at Y248 (pPD-1 Y223F), singly phosphorylated PD-1 at Y223 (pPD-1 Y248F), and non-phosphorylated PD-1 (PD-1).

We next employed the micropipette adhesion assay^28^ to quantify the affinity between SHP2 or its t-SH2 domains and the four PD-1 isoforms (Figure 4A). By measuring binding frequencies and quantifying molecular densities on the cell or bead surfaces (Figure 4B), we calculated the corresponding affinities (see Methods). Non-phosphorylated PD-1 exhibited negligible affinity for both SHP2 and the t-SH2 domains. Single phosphorylation at Y223 (pPD-1 Y248F) increased affinity by approximately one order of magnitude, indicating that ITIM phosphorylation contributes to SHP2 binding. Single phosphorylation at Y248 (pPD-1 Y223F) led to an additional ∼10-fold increase in affinity relative to pPD-1 Y248F for both SHP2 and the t-SH2 domains (Figure 4C).

**Figure 4.**
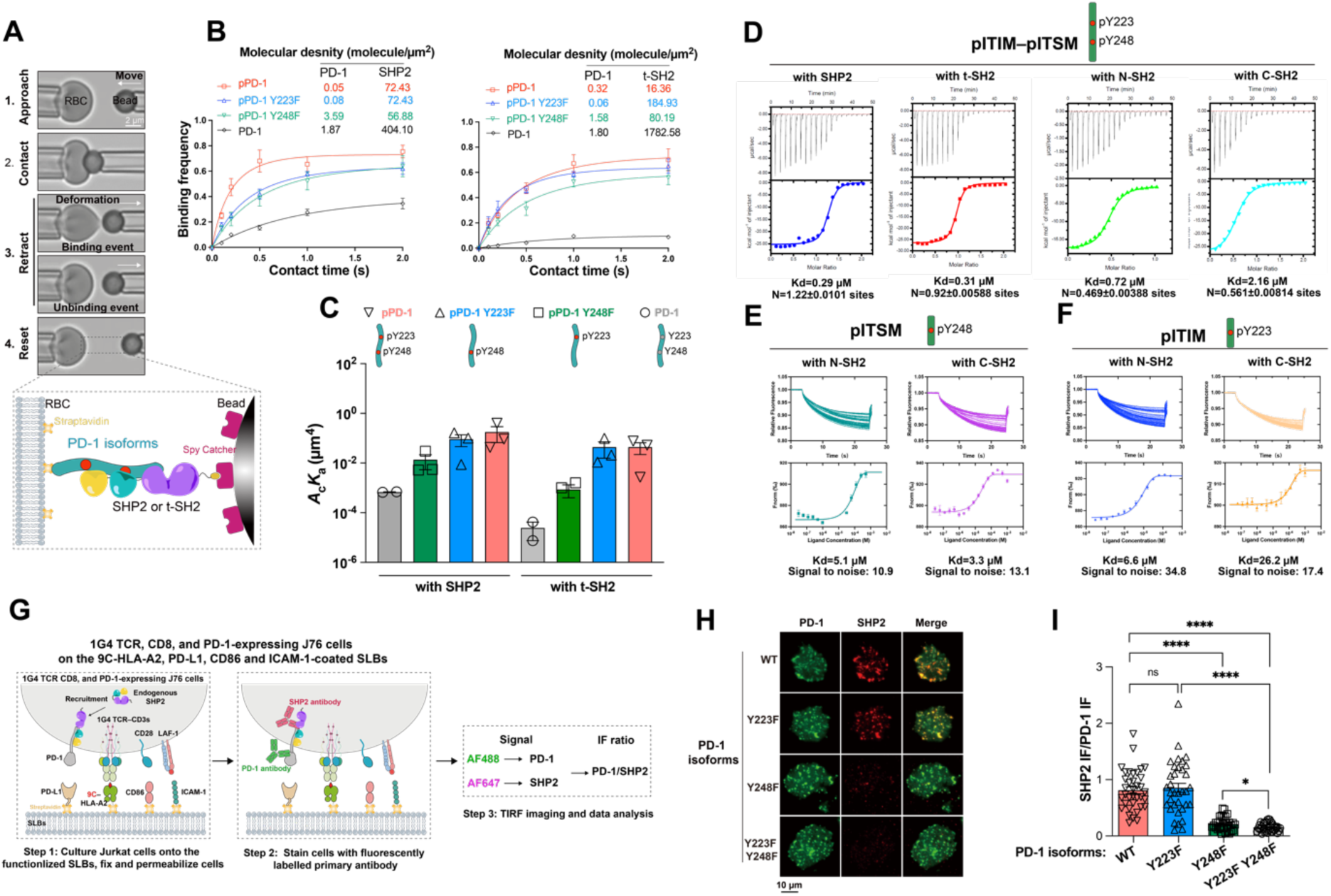
The phosphorylated sites on PD-1 precisely control the affinity and membrane recruitment of SHP2. (A) Using the micropipette adhesion frequency assay to measure the affinity between PD-1 isoforms and SHP2 isoforms. (B and C) The measured binding frequency (B) and affinity (C) between PD-1 isoforms and SHP2 isoforms using the micropipette adhesion frequency assay (*N* = 2 or 3, *n* = 2 or 3). (D) The measured affinity between PD-1 pITIM-pYITSM motif and SHP2 isoforms using the ITC assay. (E) The measured affinity between PD-1 pITSM motif and SHP2 isoforms using the MST assay. (F) The measured affinity between PD-1 pITIM motif and SHP2 isoforms using the MST assay. (G–I) The diagram (G), the representative images (H), and the results (I) of investigating the SHP2 recruitment to PD-1 using the TIRF imaging with cells interacting with the protein-functionalized SLBs (*N* = 3, *n* = 31–37). SHP2 shown in (A–C) is core SHP2 (residues: 1–525). SHP2 shown in (D) is core SHP2 (residues: 1–525) carrying the C459S mutation. PD-1 shown in (A–C) is the PD-1 intracellular region (residues: 192–288). PD-1 shown in (D) is the pITIM–pITSM motif (residues: 218–256, sequence: FSVDpYGELDFQWREKTPEPPVPCVPEQTEpYATIVFPS). PD-1 shown in (E) is the pITSM motif (residues: 242–256, sequence: VPEQTEpYATIVFPSG). PD-1 shown in (F) is the pITIM motif (residues: 218–234, sequence: VFSVDpYGELDFQWREKT). Residues of N-SH2 shown in (E–F) are 4– 103. Residues of C-SH2 shown in (E–F) are 106–218. Data in (B, C, E, and F) were presented as mean ± SD, and data in (I) were presented as mean ± SEM. The significance of the difference was determined by an unpaired two-tailed Student’s *t*-test: \*\*\*\**P* ≤ 0.0001, \**P* ≤ 0.05, and ns represents no significance. See also Figures 6–S8.

Notably, the fully phosphorylated PD-1 (pPD-1) exhibited the highest affinity for SHP2 and the t-SH2 domains; however, this affinity was not substantially higher than that of pPD-1 Y223F (Figure 4C). Together, these results demonstrate that phosphorylation at both Y223 (ITIM) and Y248 (ITSM) enhances PD-1–SHP2 binding, with ITSM phosphorylation playing the dominant role in determining binding affinity, as reflected by the similar affinities of pPD-1 and pPD-1 Y223F.

Previous studies have shown that PD-1 pITIM binds to the N-SH2 domain of SHP2, whereas PD-1 pITSM binds to either the N-SH2 or C-SH2 domain.^20^ To further define the molecular details of the interactions between SHP2 domains and PD-1 motifs, we employed isothermal titration calorimetry (ITC) to quantify the binding affinities between the fully phosphorylated pITIM–pITSM motif and full-length SHP2, the t-SH2 domains, or the isolated N-SH2 or C-SH2 domain. We found that the pITIM–pITSM motif binds with very high affinity to both full-length SHP2 and the t-SH2 domains. A similarly high affinity was observed for the isolated N-SH2 domain, but not for the C-SH2 domain (Figure 4D). These results indicate that the N-SH2 domain can engage both pITIM and pITSM motifs when sufficient N-SH2 domains are available. This observation supports a potential cellular model in which two SHP2 molecules bind to a single PD-1 molecule via their respective N-SH2 domains, engaging the pITIM and pITSM motifs.

Microscale thermophoresis (MST) analysis further revealed distinct binding profiles for the two motifs. The pITSM motif alone binds with similarly high affinity to either the N-SH2 or C-SH2 domain (Figure 4E). However, when we assessed pITIM binding using MST, we revealed relatively high affinity for the N-SH2 domain but substantially lower affinity for the C-SH2 domain (Figure 4F). Together, these findings suggest that the two SH2 domains of SHP2 can each bind a pITSM motif from separate PD-1 molecules, consistent with a previously proposed model in which PD-1 dimers are stabilized through SHP2-mediated interactions.^29,30^ These complementary measurements consistently demonstrate that while pITIM contributes to SHP2 binding, pITSM plays the dominant role in determining the overall binding affinity between SHP2 and PD-1.

### PD-1 phosphorylated states control the recruitment of SHP2 to PD-1

Having established that PD-1 phosphorylation states determine its binding affinity for SHP2, we next hypothesized that this regulated affinity governs the differential recruitment of SHP2 from the cytoplasm to PD-1 at the cell surface. To test this, we expressed 1G4 TCR, CD8, together with different PD-1 isoforms into Jurkat 76 T cells, which lack endogenous alpha and beta TCR chains (Figure S7). We incubated these cells on the supported lipid bilayers (SLBs)^31,32^ functionalized with 9C–HLA-A2, PD-L1, CD86, and ICAM- 1. Following immunostaining of transduced PD-1 and endogenous SHP2, we used total internal reflection fluorescence microscopy (TIRFM) to quantify PD-1 and SHP2 signal intensities and assess SHP2 recruitment to PD-1 across different PD-1 phosphorylation states (Figure 4G).

Wild-type (WT) PD-1 and the Y223F mutant both recruited substantial and comparable levels of SHP2, with strong colocalization between PD-1 and SHP2. In contrast, mutation of Y248 markedly reduced SHP2 recruitment. Notably, the double Y223F/Y248F mutant exhibited minimal SHP2 recruitment to PD-1 (Figures 4H and 4I). In agreement with the affinity measurements, these findings reinforce that although ITIM phosphorylation facilitates SHP2 recruitment, ITSM phosphorylation is the primary determinant of SHP2 recruitment to PD-1 at the cell surface.

### Phosphorylated PD-1 remodels SHP2 catalytic dynamics

We have thus far demonstrated that distinct PD-1 phosphorylation states remodel SHP2 conformational dynamics, binding affinity, and membrane recruitment. We next sought to determine how PD-1 phosphorylation modulates SHP2’s catalytic activity. Because SHP2’s biological functions are widely understood to depend on its phosphatase activity, we quantified SHP2 catalytic kinetics using the fluorogenic substrate 6,8-difluoro-4-methylumbelliferyl phosphate (DiFMUP).^33^ In this assay, SHP2-mediated dephosphorylation of DiFMUP generates the fluorescent product DiFMU, allowing real-time monitoring of enzymatic activity.

We first confirmed that SHP2 alone dephosphorylates DiFMUP at a low basal rate, as indicated by linear reaction curves, and that SHP2 catalytic activity scales with enzyme concentration (Figure S8A). We then introduced PD-1 motifs into the system. Using 10 nM SHP2 and 20 µM DiFMUP, we quantified DiFMU fluorescence at the 5-minute time point. The non-phosphorylated ITIM–ITSM motif had no detectable effect on SHP2 activity, whereas both singly phosphorylated motifs, pITIM–ITSM and ITIM–pITSM, significantly enhanced catalytic activity, with ITIM–pITSM producing a stronger effect. As anticipated, the fully phosphorylated pITIM–pITSM motif elicited a rapid and robust activation of SHP2 phosphatase activity (Figure S8B). As a control, the catalytically inactive SHP2 C459S mutant exhibited no detectable activity even in the presence of both phosphorylated PD-1 motifs, pITIM and pITSM (Figure S8B).

In parallel, we measured SHP2 enzymatic velocities using 10 nM SHP2 and 1 µM PD-1 motif and derived kinetic parameters at the 5-minute time point (Figure S8C). Michaelis–Menten fitting yielded V_max_, K_m_, K_cat_, and K_cat_/K_m_ values in the presence or absence of phosphorylated PD-1 motifs. Consistent with the endpoint measurements, the fully phosphorylated pITIM–pITSM motif was the most potent enhancer of SHP2 enzymatic efficiency, followed by the singly phosphorylated ITIM–pITSM and pITIM–ITSM motifs. In contrast, the non-phosphorylated ITIM–ITSM motif had minimal impact on SHP2 catalytic kinetics (Figure S8D).

### pPD-1 drives SHP2’s non-catalytic regions to dissolve pTCR and pCD28 condensates

Although our preceding experiments establish that PD-1 phosphorylation activates SHP2 by promoting its closed-to-open conformational transition and enhancing phosphatase activity, these findings do not fully account for how PD-1 suppresses higher-order organization of TCR signaling assemblies, nor for the pronounced temporal lag between SHP2 recruitment and downstream TCR dephosphorylation.^15^

Recent studies have shown that SHP2 can undergo liquid–liquid phase separation (LLPS) through multivalent electrostatic interactions involving its PTP domains, and that disease-associated mutants stabilized in the open conformation further potentiate this behavior.^34^ These observations raise the possibility that PD-1-driven SHP2 activation may suppress TCR signaling not only through catalytic dephosphorylation, but also via biophysical mechanisms that remodel signaling condensates.

Although CD28 has been identified as a primary substrate of the PD-1–SHP2 signalosome,^14,35^ components of the TCR complex are also well-established SHP2 targets.^15,36,37^ Moreover, SHP2 has been reported to form condensates in a phosphatase-dependent manner, potentially amplifying its inhibitory function in T cells. ^34^ Building on our previous work defining LLPS mechanisms of TCR/Lck^38^ and CD28/Lck^39^ assemblies, we next asked how the PD-1–SHP2 axis influences the formation and stability of these condensates.

To this end, we first reconstituted phosphorylated CD28 (pCD28)/Lck condensates on supported lipid bilayers (SLBs) in the presence of PD-1 with defined phosphorylation states. Strikingly, PD-1 was efficiently incorporated into pCD28/Lck condensates in a phosphorylation-dependent manner, with increased PD-1 phosphorylation promoting its partitioning into these assemblies (Figure 5A). This suggests that PD-1 phosphorylation licenses its entry into CD28 signaling condensates.

**Figure 5.**
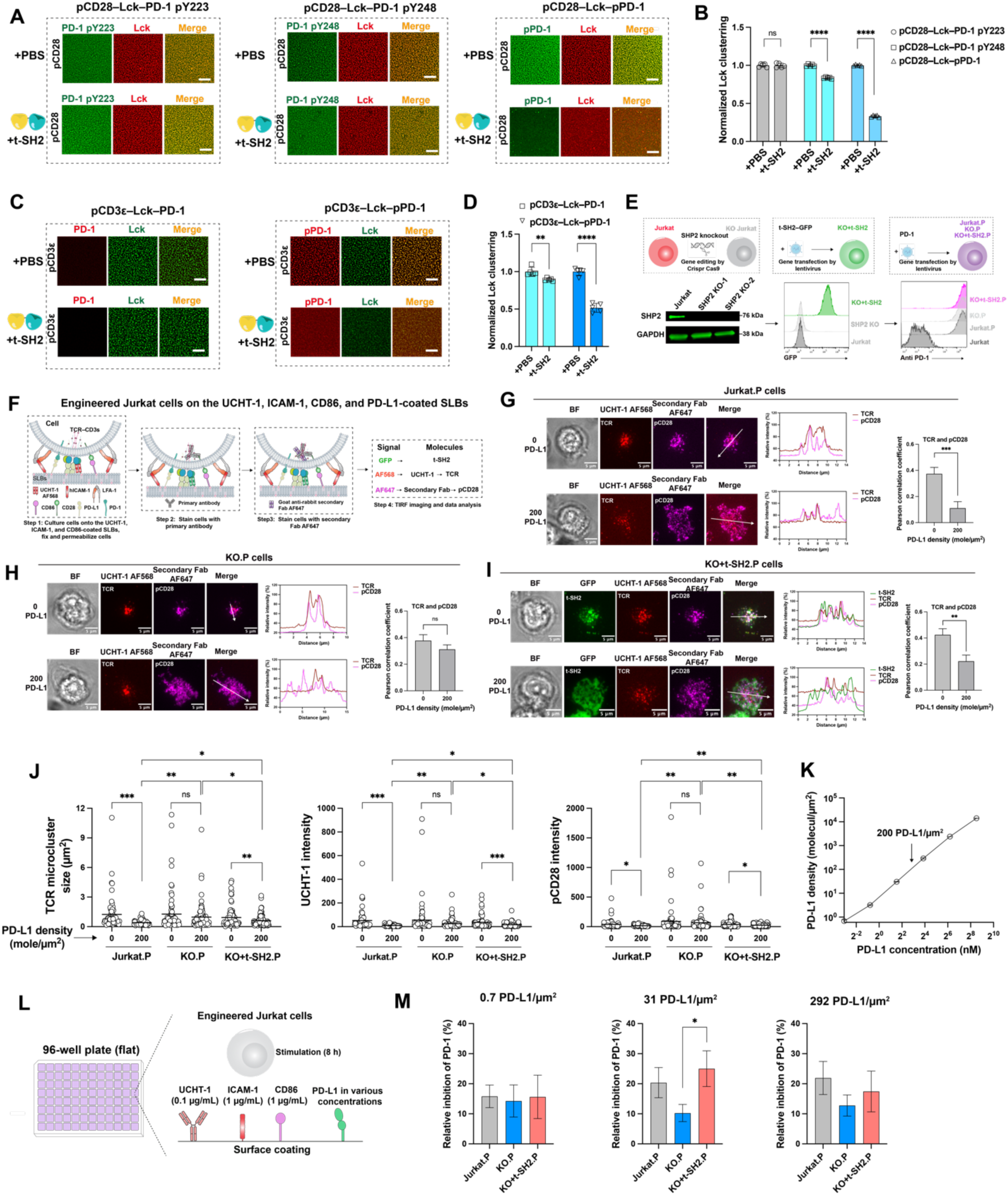
PD-1 drives SHP2’s non-catalytic inhibition of Jurkat T cell activation through disrupting pTCR and pCD28 proximal signaling clusters. (A) Representative confocal images of liquid–liquid phase separation (LLPS) formed by Lck–pCD28–phosphorylated PD-1 with the addition of t-SH2 domains of SHP2 or PBS. The more phosphorylation of PD-1, the more integration of PD-1 into LLPS. (B) The quantified Lck clustering in LLPS with t-SH2 or PBS. t-SH2 domains significantly disrupt the LLPS depending on the phosphorylation of PD-1 (*N* = 3, *n* = 5). (C) Representative confocal images of LLPS formed by Lck–pCD3ε–pPD-1 with the addition of t-SH2 domains of SHP2 or PBS. The more phosphorylation of PD-1, the more integration of PD-1 into LLPS. (D) The quantified Lck clustering in LLPS with t-SH2 or PBS. t-SH2 domains significantly disrupt the LLPS depending on the phosphorylation of PD-1 (*N* = 3, *n* = 5). (E) The generation of the engineered Jurkat T cells. (F) The processes of applying the QuEST method to investigate the TCR, CD28 signaling, which is regulated by the PD-1–PD-L1 signaling in different engineered Jurkat T cells. (G and H) Representative TIRF images of TCR clustering, pCD28 clustering, and colocalization analysis in Jurkat (G) and SHP2 KO Jurkat T cells (H) (*N* = 3, *n* = 31–46). (I) Representative TIRF images of TCR clustering, pCD28 clustering, t-SH2 clustering, and colocalization analysis in SHP2 KO+t-SH2 Jurkat T cells. (*N* = 3, *n* = 32–60). (J) The quantified size of TCR microclusters, TCR intensity, and pCD28 intensity from the experiment of (G–I) (*N* = 3, *n* = 54–92). (K) The quantified PD-L1 density on the coverslip with various coating concentrations using the QuEST method (*N* = 3, *n* = 3). (L) The processes of investigating SHP2’s non-catalytic inhibition of Jurkat T cell activation in cells. (M) The relative PD-1 inhibition of engineered Jurkat T cells with 0.7, 31, and 292 PD-L1/µm^2^ on the surface, respectively. The relative PD-1 inhibition values were calculated by subtracting the data shown in Figure S12 from 100% (*N* = 3, *n* = 6–8). pCD3ε shown here is the intracellular region (residues: 153–207). PD-1 shown here is the intracellular region (residues: 192–288). CD28 shown here is the intracellular region (residues: 180–220). pPD-1 was generated through the phosphorylation of PD-1 by the kinase domain of Lck. pCD28 and pCD3ε were commercially synthesized. Lck shown here is the region containing the Unique, SH3, and SH2 domains. t-SH2 domains contain residues: 1–220. Data in (B, D, and K) were presented as mean ± SD, data in (G–I and M) were presented as mean ± SEM. Scale bar size in (A and C) is 10 µm. The significance of the difference was determined by an unpaired two-tailed Student’s *t*-test: \*\*\*\**P* ≤ 0.0001, \*\*\**P* ≤ 0.001, \*\**P* ≤ 0.01, \**P* ≤ 0.05, and ns represents no significance. See also Figures 9–S13.

We next examined how the non-catalytic regions of SHP2 regulate pCD28/Lck condensates. Unexpectedly, the tandem SH2 (t-SH2) domains of SHP2 potently disassembled these condensates in a PD-1 phosphorylation-dependent manner. Fully phosphorylated PD-1 (pPD-1) exerted the strongest effect, followed by PD-1 pY248, whereas PD-1 pY223 failed to promote condensate dissolution (Figures 5A and 5B). These findings further underscore the critical role of the PD-1 ITSM motif in mediating PD-1 signaling. We then extended these observations to TCR condensates by reconstituting phosphorylated CD3ε (pCD3ε)/Lck assemblies on SLBs in the presence of either unphosphorylated or fully phosphorylated PD- 1. Consistent with the CD28 system, PD-1 was incorporated into pCD3ε/Lck condensates only upon full phosphorylation (Figure 5C). Addition of SHP2 t-SH2 domains induced robust dissolution of these condensates, and this effect strictly depended on PD-1 phosphorylation (Figures 5C and 5D). These results demonstrate that recruitment of SHP2 t-SH2 domains is sufficient to disrupt TCR/Lck condensates, revealing a regulatory mechanism that does not inherently require SHP2 catalytic activity.

To further support a non-catalytic mode of inhibition, we repeated these experiments using a phosphatase-dead SHP2 mutant (C459S). In the presence of phosphorylated PD-1, SHP2 C459S disrupted TCR/Lck condensates to a similar extent as the isolated t-SH2 domains (Figures 9A and 9B), reinforcing the conclusion that SHP2-mediated suppression can occur independently of its enzymatic activity.

Importantly, these experiments (Figures 5A–5D and S9) were performed at low protein concentrations on SLBs that preclude SHP2 self-phase separation, thereby excluding contributions from SHP2 LLPS. To validate this, we conducted three-dimensional phase separation assays at higher protein concentrations. Under these conditions, SHP2 C459S formed phase-separated droplets that efficiently recruited phosphorylated PD-1 (Figure S10A), confirming their direct interaction. The same protein, but at low concentrations on the SLBs, did not form any condensates, no matter the phosphorylation of PD-1 (Figure S10B).

Together, these findings indicate that PD-1-recruited SHP2, via its SH2 domains, could suppress TCR/Lck and CD28/Lck condensates through a non-catalytic mechanism, independently of phosphatase activity.

### PD-1 drives SHP2’s non-catalytic inhibition of Jurkat T cell activation through disrupting TCR and pCD28 proximal signaling clustering

The non-catalytic inhibition of TCR and CD28 condensates by the SHP2 t-SH2 domains, as revealed by our LLPS reconstitution assays, together with our recent finding that activated PD-1 is highly enriched within TCR clusters,^40^ prompted us to investigate whether this mechanism operates in T cells to modulate activation. To this end, we first generated SHP2 knockout (KO) Jurkat T cells and reconstituted them with t-SH2–GFP together with PD-1 (Figure 5E).

Using the QuEST method recently developed by us,^40^ we stimulated these engineered cells on supported lipid bilayers (SLBs) functionalized with UCHT-1 Fab (36 molecules/µm^2^), ICAM-1 ectodomain (177 molecules/µm^2^), CD86 ectodomain (200 molecules/µm^2^), and either in the presence or absence of PD-L1 (200 molecules/µm^2^). We then quantified fluorescence signals corresponding to t-SH2 (GFP), TCR (UCHT-1), and CD28 phosphorylation (pCD28) (Figure 5F).

In control Jurkat T cells, TCR and pCD28 formed prominent microclusters in the absence of PD-1 signaling, whereas the presence of PD-L1 markedly reduced these assemblies and their colocalization (Figure 5G). Quantitative analysis revealed significant decreases in TCR microcluster size, TCR signal intensity, and pCD28 phosphorylation upon PD-1 engagement (Figure 5J). In contrast, these inhibitory effects were abolished in SHP2 KO cells, where PD-1 signaling failed to reduce microcluster size, colocalization, or signaling strength (Figures 5H and 5J), consistent with the requirement for SHP2 in PD-1-mediated suppression.

Remarkably, reintroduction of the isolated t-SH2 domains into SHP2 KO cells restored inhibition: t-SH2 expression significantly disrupted both TCR and pCD28 proximal signaling clusters and colocalization (Figures 5I and 5J). Although the magnitude of this effect was somewhat weaker than that observed in wild-type Jurkat cells, these results demonstrate that SHP2 t-SH2 domains alone are sufficient to mediate suppression of signaling condensates in cells.

We next examined how this non-catalytic mechanism influences downstream T cell activation. Using QuEST,^40^ we first calibrated PD-L1 surface density to match ∼200 molecules/µm^2^, a physiologically relevant level consistent with our SLB assays^41^ (Figures 5K and S11). Jurkat, SHP2 KO, and KO+t-SH2 cells were then stimulated on surfaces presenting UCHT-1, ICAM-1, CD86, and varying densities of PD-L1 (Figure 5L).

At low PD-L1 densities, Jurkat cells exhibited significantly reduced CD69 expression compared to SHP2 KO cells, indicating that SHP2 is required for efficient PD-1-mediated inhibition under these conditions. At higher PD-L1 densities, however, CD69 expression was strongly suppressed in both cell types (Figure S12), likely reflecting compensatory contributions from SHP1 in the absence of SHP2.

Notably, KO+t-SH2 cells recapitulated the reduced CD69 expression observed in Jurkat cells at low PD-L1 densities, consistent with a non-catalytic inhibitory role of the SHP2 SH2 domains. Unexpectedly, at high PD-L1 densities, KO+t-SH2 cells failed to suppress activation to the same extent as Jurkat cells, suggesting that t-SH2 occupancy of PD-1 may competitively limit recruitment of SHP1 under these conditions (Figure S12). These findings may support a model that t-SH2 of SHP2 inhibits T cell activation in a two-stage pattern.

Finally, at PD-L1 densities of 31 and 292 molecules/µm^2^, which encompass the physiologically relevant range consistent with our SLB assays (Figures 5F–5J), the KO+t-SH2 cells exhibited stronger PD-1-mediated inhibition than SHP2 KO cells, whereas this difference was lost at very low PD-L1 densities (Figure 5M).

To further validate the non-catalytic mechanism, we generated SHP2 KO cells reconstituted with a phosphatase-dead mutant (C459S) (Figure S13A). These cells displayed PD-1-dependent CD69 suppression comparable to KO+t-SH2 cells (Figure S13B), confirming that SHP2 can inhibit T cell activation independently of its catalytic activity.

## Discussion

In this study, we establish a multiscale framework to elucidate how PD-1 regulates SHP2 structural and functional dynamics to suppress T cell activation. Our atomic structural analysis and single-molecule biophysical analyses demonstrate that binding of phosphorylated PD-1 reshapes the conformational dynamics and energy landscape of SHP2 at single-molecule resolution, accelerating SHP2’s closed-to-open transition by at least 94-fold. Other than conventional catalytic function, the opened SHP2 conformation, especially its t-SH2 domains, performs as a biophysical disruptor to rapidly disassemble pTCR and pCD28 condensates to inhibit T cell signaling.

This structural transition provides the basis for a dual-layered mechanism of T cell suppression. While PD-1 phosphorylation enhances SHP2 phosphatase activity (the catalytic layer), we identify a previously unrecognized non-catalytic mechanism mediated by the highly released t-SH2 domains. In the reconstituted SLB system, the t-SH2 domains physically dismantle pTCR/Lck- and pCD28/Lck-formed condensates in a strictly PD-1 phosphorylation-dependent manner, independent of SHP2 catalytic activity. Our data also clarify the role of phase separation in this system. We and others^34^ have proved that while SHP2 is capable of forming condensates at high concentrations, the inhibitory effects we observe on supported membranes occur under conditions where SHP2 does not phase separate. Thus, SHP2-mediated suppression of TCR and CD28 condensates is primarily driven by competitive and multivalent SH2 interactions rather than SHP2’s LLPS.

Importantly, this non-catalytic mechanism was also observed in cells. Reintroduction of SHP2 t-SH2 domains into SHP2-deficient Jurkat T cells restores PD-1-dependent suppression of TCR microclusters, CD28 phosphorylation, TCR/pCD28 colocalization, and CD69 upregulation, although not to the full extent of wild-type SHP2. These results demonstrate that SH2-mediated interactions are sufficient to mediate a substantial component of PD-1 inhibitory function, while also indicating that catalytic activity contributes additional suppression under certain conditions. Although the SHP2-deficient Jurkat T cells displayed activation inhibition with high PD-L1 concentrations, this might be because PD-1 also recruits SHP1, although not as much as SHP2.^15^ Moreover, a recent study also demonstrated that PD-1 can suppress T cell signaling independent of SHP2,^42^ which, to some extent, explains the inhibition of Jurkat T cell activation in SHP2 KO cells with high PD-L1 concentrations.

Placing these findings into a broader biological context, the highly open SHP2 acts as a competitive scaffold or spatial steric hindrance. This dual-layered model resolves the temporal paradox between rapid SHP2 recruitment and delayed substrate dephosphorylation, that is, the physical disassembly of signaling condensates provides an immediate, coarse blockade of immune synapse assembly, which is subsequently reinforced by the sustained biochemical dephosphorylation of signaling substrates. We recently found that the stoichiometry of ZAP-70 to TCR remains as low as 1:1 throughout T cell activation,^40^ suggesting limited ZAP-70 occupancy at the receptor complex. Given that ZAP-70 is also a t-SH2 domain-containing protein capable of engaging the TCR/CD28 signaling network, these findings raise the possibility that SHP2-mediated non-catalytic inhibition competes with ZAP-70 recruitment, thereby suppressing proximal TCR/CD28 signal transduction.

Our structural analysis shows that binding of phosphorylated PD-1 stabilizes SHP2 in a fully open conformation that is more extensively rearranged than previously described open states, including disease-associated mutants.^19^ This conformation is characterized by large-scale domain reorganization. Complementing this structural snapshot, single-molecule magnetic tweezers experiments reveal that SHP2 intrinsically possesses closed and open conformations. Importantly, PD-1 phosphorylation dramatically accelerates this transition by at least 94-fold, indicating that ligand engagement primarily acts by reshaping the conformational energy landscape of SHP2 to favor the open state rather than simply inducing a static structural switch.

Our quantitative binding and cellular recruitment data further reveal a clear hierarchy between the two PD-1 tyrosine motifs. While phosphorylation at Y223 (ITIM) contributes to SHP2 interaction, phosphorylation at Y248 (ITSM) is the dominant determinant of binding affinity and membrane recruitment. Consistently, ITSM phosphorylation alone is sufficient to achieve near-maximal SHP2 binding, whereas ITIM phosphorylation plays a more modest, supportive role. These findings suggest that PD-1 signaling strength is encoded in its phosphorylation pattern, with ITSM acting as the primary driver of SHP2 engagement and ITIM providing an additional layer of modulation.

Together, our findings support a model in which PD-1 inhibits T cell activation through two coordinated but mechanistically distinct layers. First, PD-1 phosphorylation promotes SHP2 opening and enhances its phosphatase activity, enabling enzymatic dephosphorylation of signaling substrates. Second, and more immediately, PD-1-recruited SHP2 disrupts pTCR and pCD28 signaling assemblies through multivalent SH2–phosphotyrosine interactions that destabilize condensates. This non-catalytic mechanism provides a direct explanation for the previously observed temporal gap between rapid SHP2 recruitment and delayed substrate dephosphorylation,^15^ as structural disruption of signaling clusters can occur independently of enzymatic activity.

These mechanistic insights have important implications for therapeutic targeting of the PD-1/SHP2 axis. Current immune checkpoint therapies block PD-1/PD-L1 interactions and thereby prevent SHP2 recruitment. However, our results suggest that SHP2 function can also be modulated at the level of conformational dynamics or SH2-mediated interactions. For example, allosteric inhibitors such as SHP099 stabilize the closed conformation and suppress dynamic switching, while disruption of SH2–phosphotyrosine interactions could selectively impair the non-catalytic inhibitory mechanism. Targeting these distinct layers may provide alternative or complementary strategies to modulate T cell responses.

In summary, we show that PD-1 phosphorylation controls SHP2 activity through coordinated regulation of conformational dynamics, binding affinity, and both catalytic and non-catalytic functions. By demonstrating that SHP2 can suppress T cell signaling independently of its phosphatase activity, our study expands the current view of immune checkpoint signaling and highlights the importance of structural and mesoscale mechanisms in regulating T cell activation.

### Limitations of the study

Despite the breadth of approaches used in this study, several limitations should be considered. First, the crystal structure was determined using PD-1-derived phosphorylated motifs rather than the full-length PD-1 intracellular tail, which may exhibit additional conformational flexibility or regulatory interactions in vivo. Second, magnetic tweezers experiments involve externally applied forces that, while physiologically relevant in magnitude, do not fully recapitulate the complex mechanical environment of the immunological synapse. Finally, phase separation assays were conducted in simplified in vitro systems and may not fully capture the regulatory constraints present in cells.

## Supporting information

STAR Methods

Validation of the structure

## Acknowledgments

We want to express our gratitude to Dr. Hu Chen at Xiamen University, Dr. Shimin Le, Dr. Mingxi Yao, and Dr. Jie Yan at the National University of Singapore, Dr. Wei Li at the Institute of Physics, Chinese Academy of Sciences, Dr. Junwei Liu and Chuxuan Xiao at Zhejiang University, for their generous help in establishing the single-molecule magnetic tweezers (MT). We thank Dr. Sanhuang Fang and Jiajia Wang from the core facilities in Zhejiang University School of Medicine for the technical assistance in TIRF imaging and cell sorting, respectively. We thank the Oxford-Zeiss Centre of Excellence for Biomedical Imaging and Flow Cytometry Facilities at the University of Oxford. We thank Elke Kurz for her help in organizing experimental reagents. We thank the beamline staff of BL19U1 at Shanghai Synchrotron Radiation Facility for their help with data collection. We thank the staff members of the Large-scale Protein Preparation System at the National Facility for Protein Science in Shanghai (NFPS), Shanghai Advanced Research Institute, Chinese Academy of Sciences, China, for providing technical support and assistance in ITC data collection. We thank Fiji for the imaging analysis. This research was financially supported by the following foundations: the National Natural Science Foundation of China (32371250 to C.Z., T2394511, T2394510, and 92359303 to W.C., 32090044, 11672317 to J.L., and T2394512 to H.C.), the National Natural Science Youth Foundation of China (12202385 to P.F., 32200549 to H.C., and 32101052 to P.W.), the China Postdoctoral Science Foundation (2022M722735 to P.F.), the Key Project of Natural Science Foundation of Zhejiang Province (LZ24C050002 to C.Z.), the National Science and Technology Major Project (2023ZD0501300 to W.C.), the Chinese Academy of Medical Sciences (CAMS) Innovation Fund for Medical Science (CIFMS), China (2024-I2M-2-001-1 to M.L.D.), the Kennedy Trust for Rheumatology Research Cell Dynamics Platform (KENN 20 21 17 to M.L.D.), and the Wellcome Trust (100262Z/12/Z and 224343/Z/21/Z to M.L.D.).

## Author contributions

W.C., C.Z., J.L., M.L.D., and P.F. provided project supervision. P.F. established the single-molecule magnetic tweezers (MT). P.F. performed MT experiments and MT data analysis. J.G., P.J., P.W., and P.F. constructed plasmids. J.G., P.J., P.W., Lu.C., and Li.C. purified recombinant protein. P.F. and J.G. constructed cell lines. P.F. and S.V. produced the supported lipid bilayers. P.F. handled the TIRF imaging. P.J. and C.Z. resolved and analyzed X-ray crystal structures. H.C. performed in vitro protein phosphorylation and identification. J.G. and P.F. measured protein affinity using the micropipette adhesion assay. P.J. conducted protein affinity measurements via isothermal titration calorimetry and microscale thermophoresis. P.J. performed SHP2 enzymatic activity measurements. H.C. carried out liquid–liquid phase separation experiments. Y.Z. performed molecular dynamics simulations. P.F. performed the cellular experiments. P.F. analyzed the overall data to generate figures and legends. P.F. wrote the primary manuscript with contributions from the other authors. P.F., J.G., M.L.D., C.Z., J.L., and W.C. revised the manuscript, and all authors approved the manuscript.

## Data availability

The atomic coordinates and structure factors have been deposited in the Research Collaboratory for Structural Bioinformatics Protein Data Bank (RCSB PDB, https://www.rcsb.org/) with the following accession codes: 21BJ.

## Declaration of generative AI and AI-assisted technologies

During the preparation of this work, the authors used ChatGPT in order to polish the written language. After using ChatGPT, the authors reviewed and edited the content as needed and take full responsibility for the content of the publication.

---

**Figure S1.**
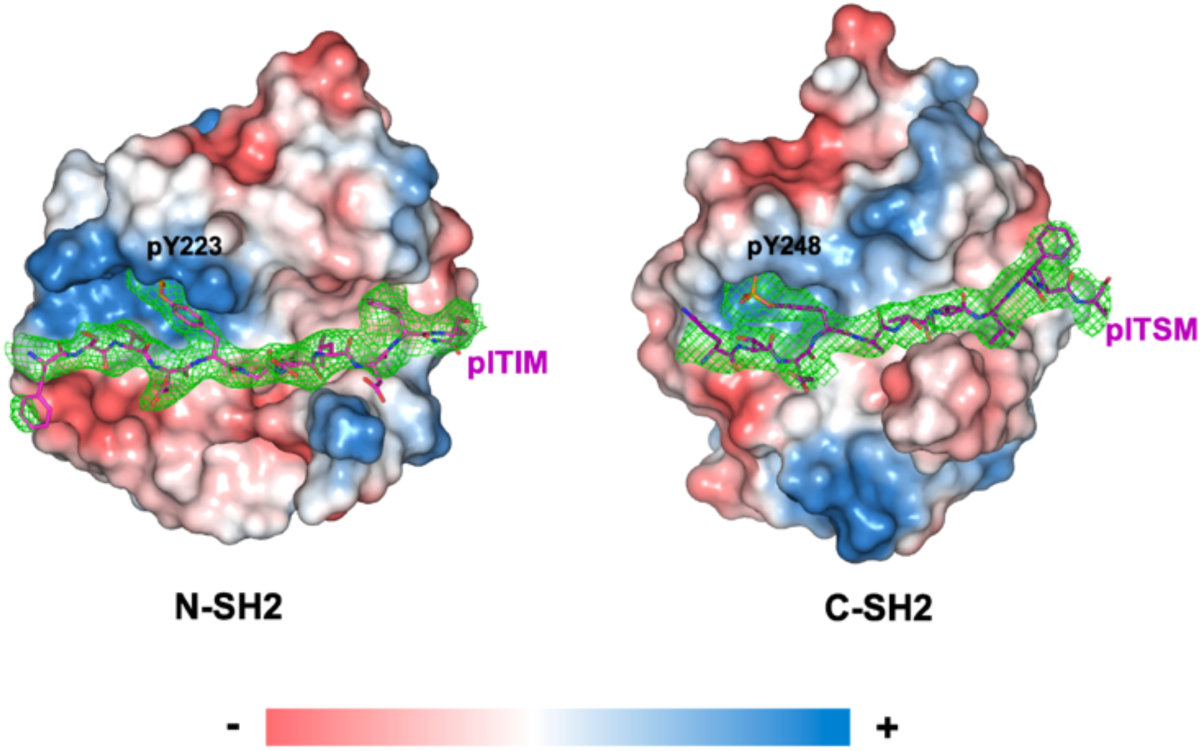
Omit maps (green mesh) of pITIM and pITSM contoured at 1.5 σ. Electrostatic surface charge of N-SH2 (left) and C-SH2 (right) domains is also displayed. Related to Figure 1.

**Figure S2.**
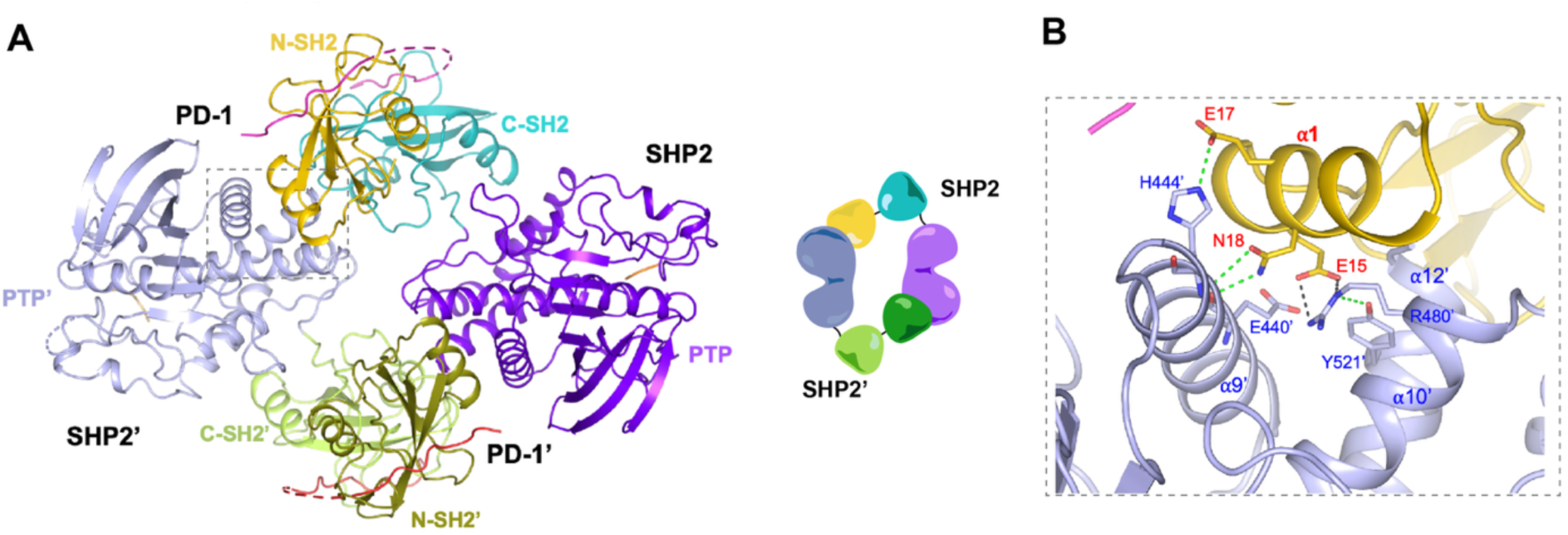
Dimeric SHP2–PD1 motif complex structure. Related to Figure 1. (A) Dimeric SHP2-PD1 motif complex structure observed in crystal. (B) Close-up view of the interface between SHP2 N-SH2 with the PTP domain from the other SHP2 molecule in the dimer. Green dashed lines indicate hydrogen bonds, grey dashed lines indicate salt bridges. SHP2 shown here is core SHP2 (residues 1–525) carrying the C459S mutation. pITIM motif (residues: 218–234, sequence: VFSVDpYGELDFQWREKT), and pITSM motif (residues: 242–256, sequence: VPEQTEpYATIVFPSG).

**Figure S3.**
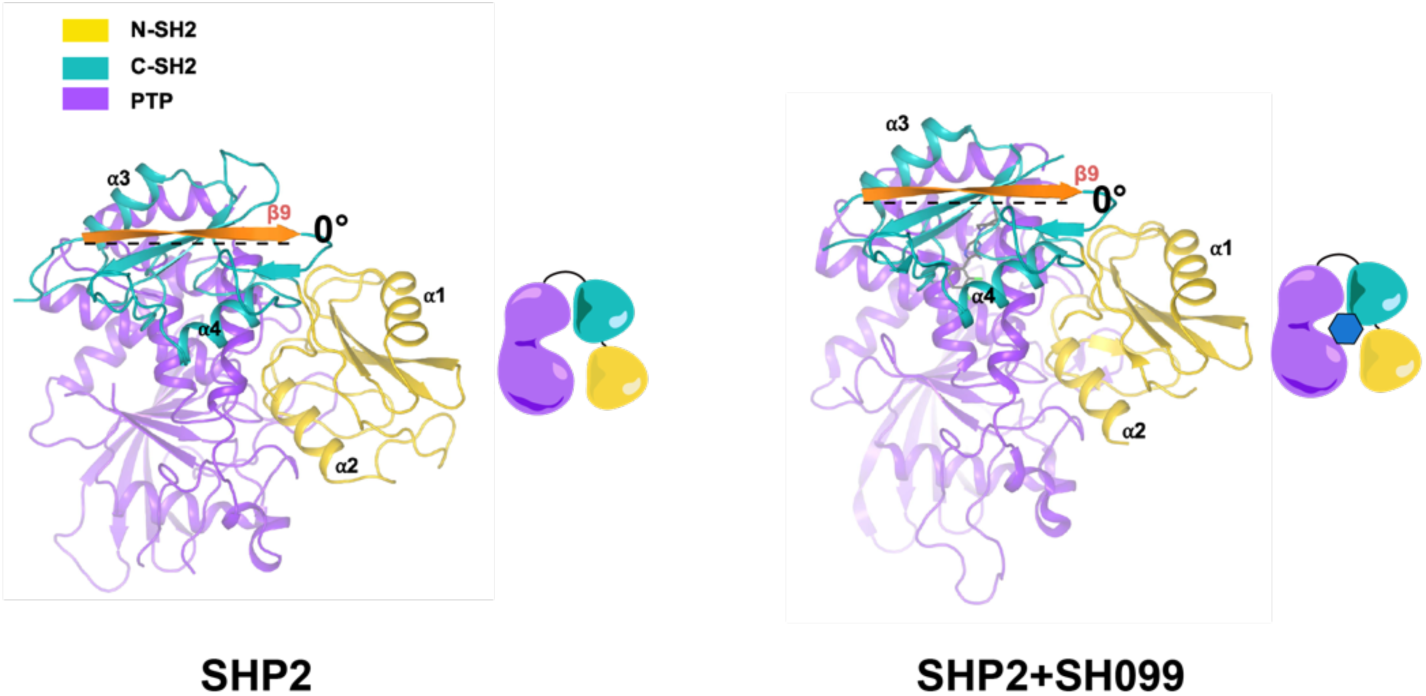
Crystal structures of SHP2 alone (PDB: 2SHP) or in complex with the allosteric inhibitor SHP099 (PDB: 5EHR). Related to Figure 3. β9 in the C-SH2 domain is colored orange to highlight the orientation of the C-SH2 domain, and the rotation degree is labelled. SHP2 shown here is core SHP2 (residues: 1–525).

**Figure S4.**
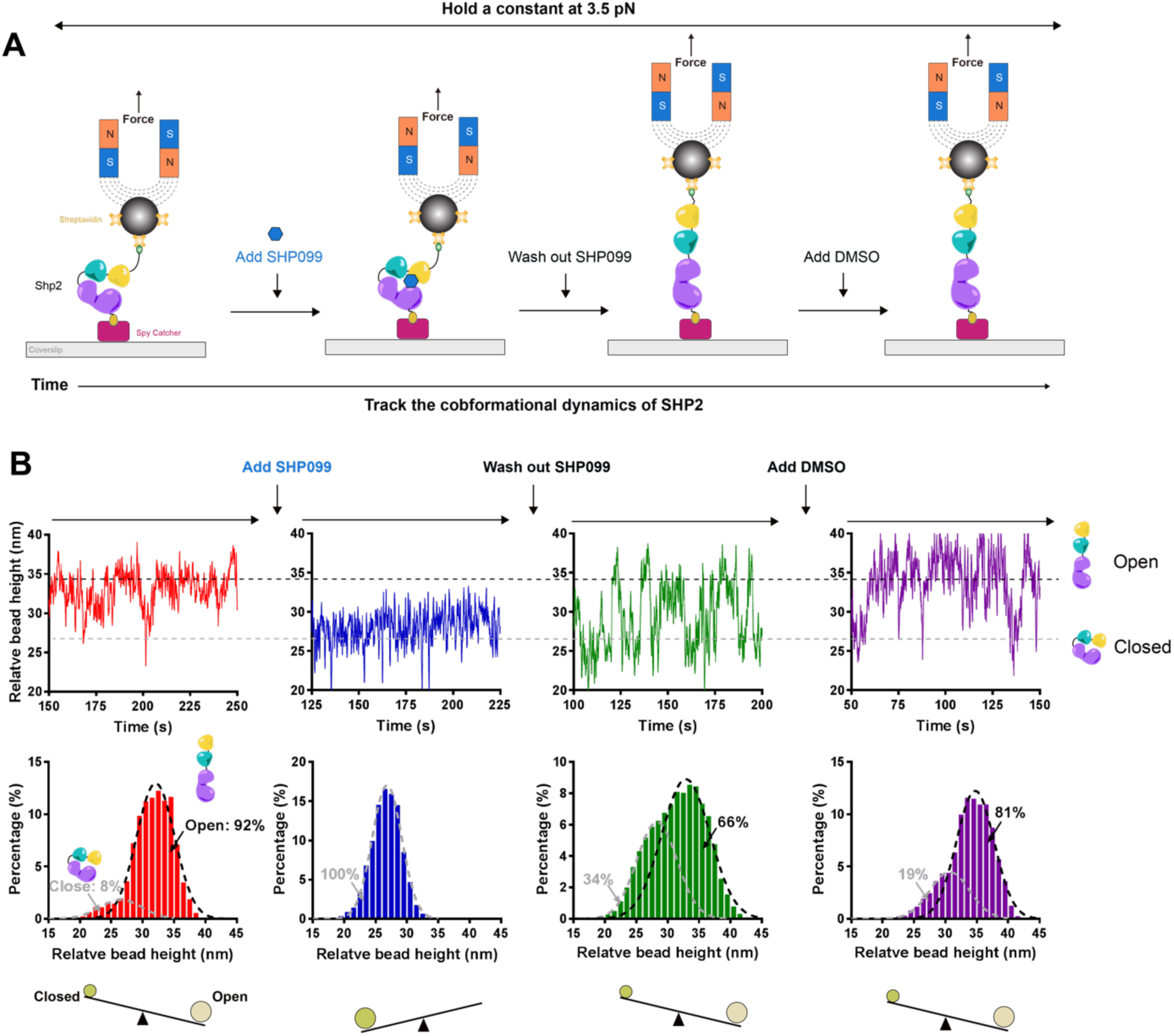
The allosteric inhibition of SHP099 on SHP2 is reversible. Related to Figure 3. (A) The diagram of using the MT to track the dynamics of conformational changes of SHP2 with or without SHP099 under 3.5 pN constant force, SHP2 with DMSO serves as the control to eliminate the effect of DMSO on the SHP2 conformations. (B) The dynamics of conformational changes of SHP2 with or without SHP099. Although the experiments were conducted continuously before and after each group, the time axis for each graph was reset, and a 100-second segment was selected for display. SHP2 shown here is core SHP2 (residues: 1–525).

**Figure S5.**
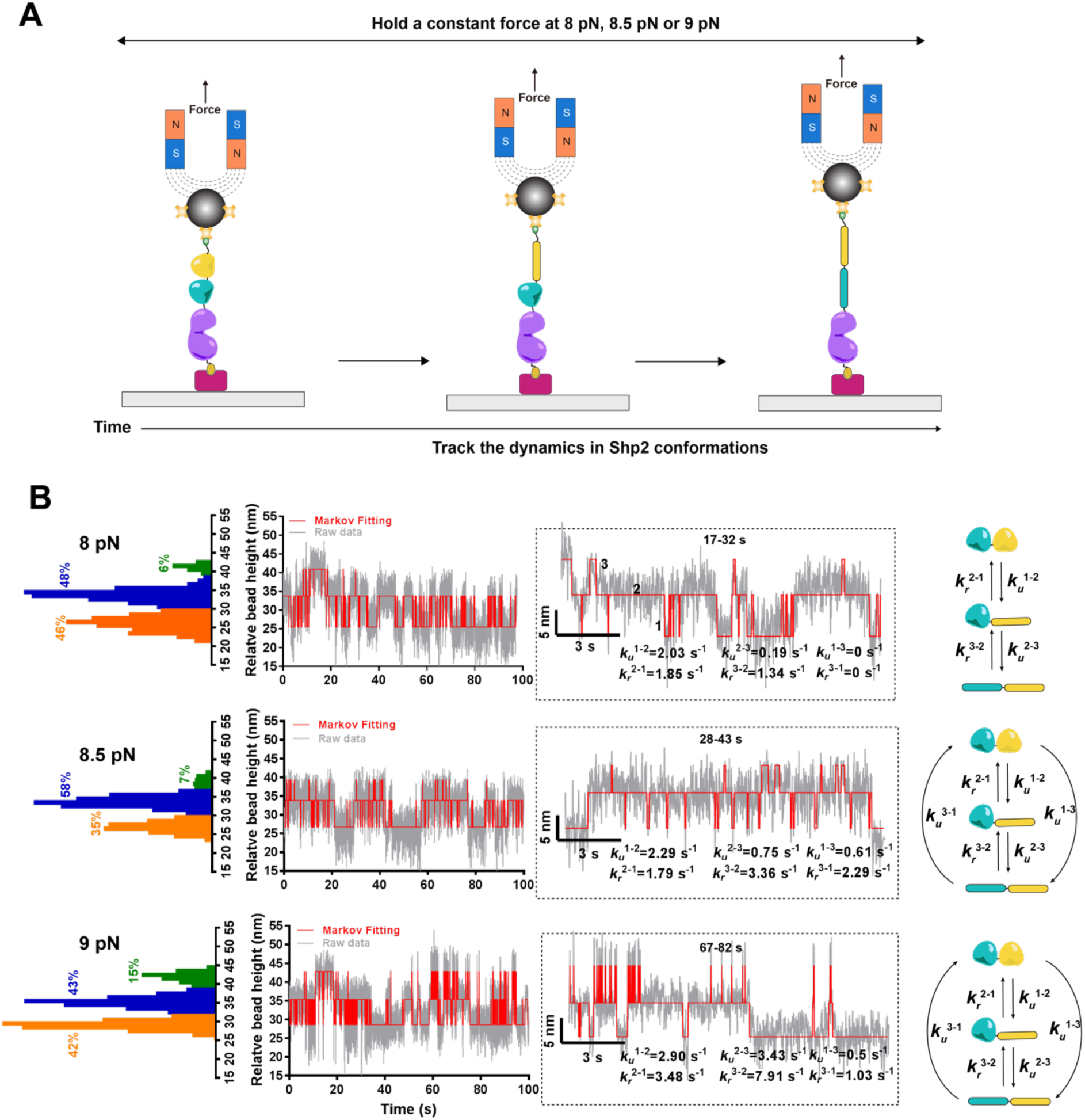
The unfolding and refolding conformational switch of the N-SH2 and C-SH2 domains in SHP2 is captured and quantified. Related to Figures 2 and 3. (A) The diagram of using the MT to track the unfolding and refolding conformational switch of the N-SH2 C-SH2 region in SHP2 under 8 pN, 8.5 pN, or 9 pN constant forces. (B) The dynamics of the unfolding and refolding conformational switch of the N-SH2 and C-SH2 domains in SHP2. Switch rates and paths between different conformational states were labelled. Although the experiments were conducted continuously before and after each group, the time axis for each graph was reset to zero, and a 100-second segment was selected for display. SHP2 shown here is core SHP2 (residues: 1–525).

**Figure S6.**
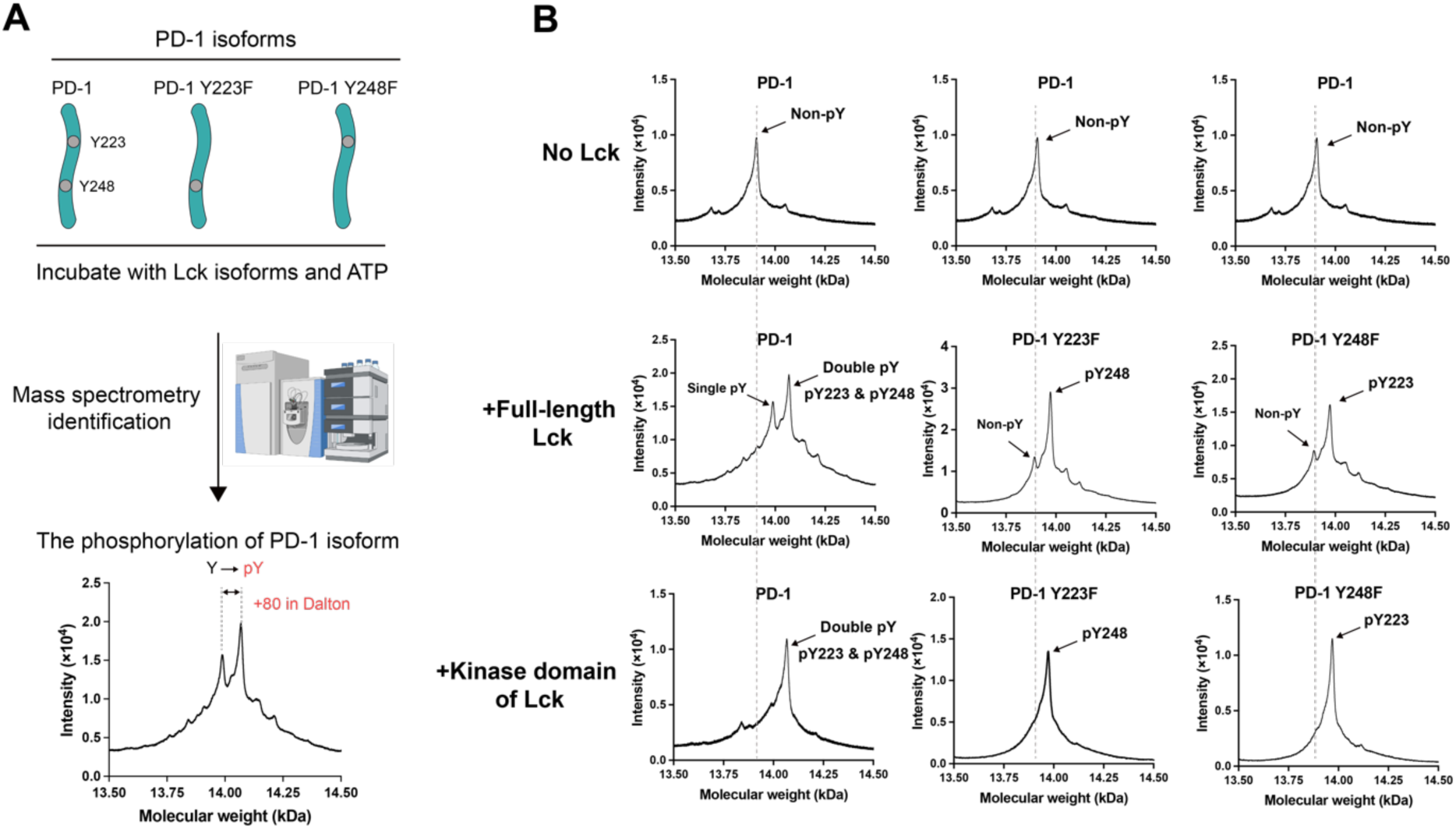
PD-1 is fully phosphorylated by the kinase domain of Lck. Related to Figure 4. (A) The diagram of using full-length Lck or the kinase domain of Lck to phosphorylate PD-1, and the subsequent phosphorylation identification by mass spectrometry. (B) Validation on the phosphorylation of PD-1. PD-1 shown here is the PD-1 intracellular region (residues: 192–288).

**Figure S7.**
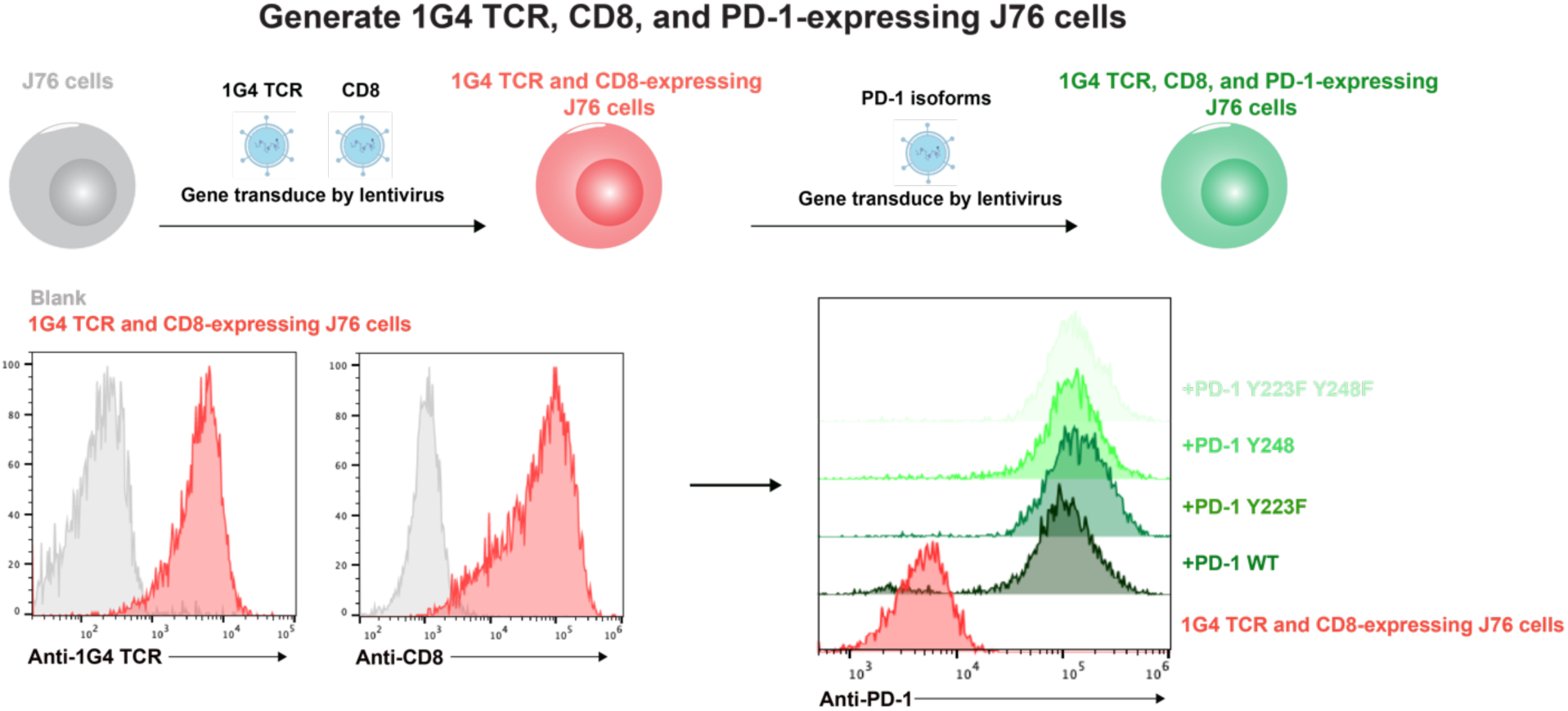
Generation of 1G4 TCR, CD8, and PD-1-expressing J76 cells. Related to Figure 4.

**Figure S8.**
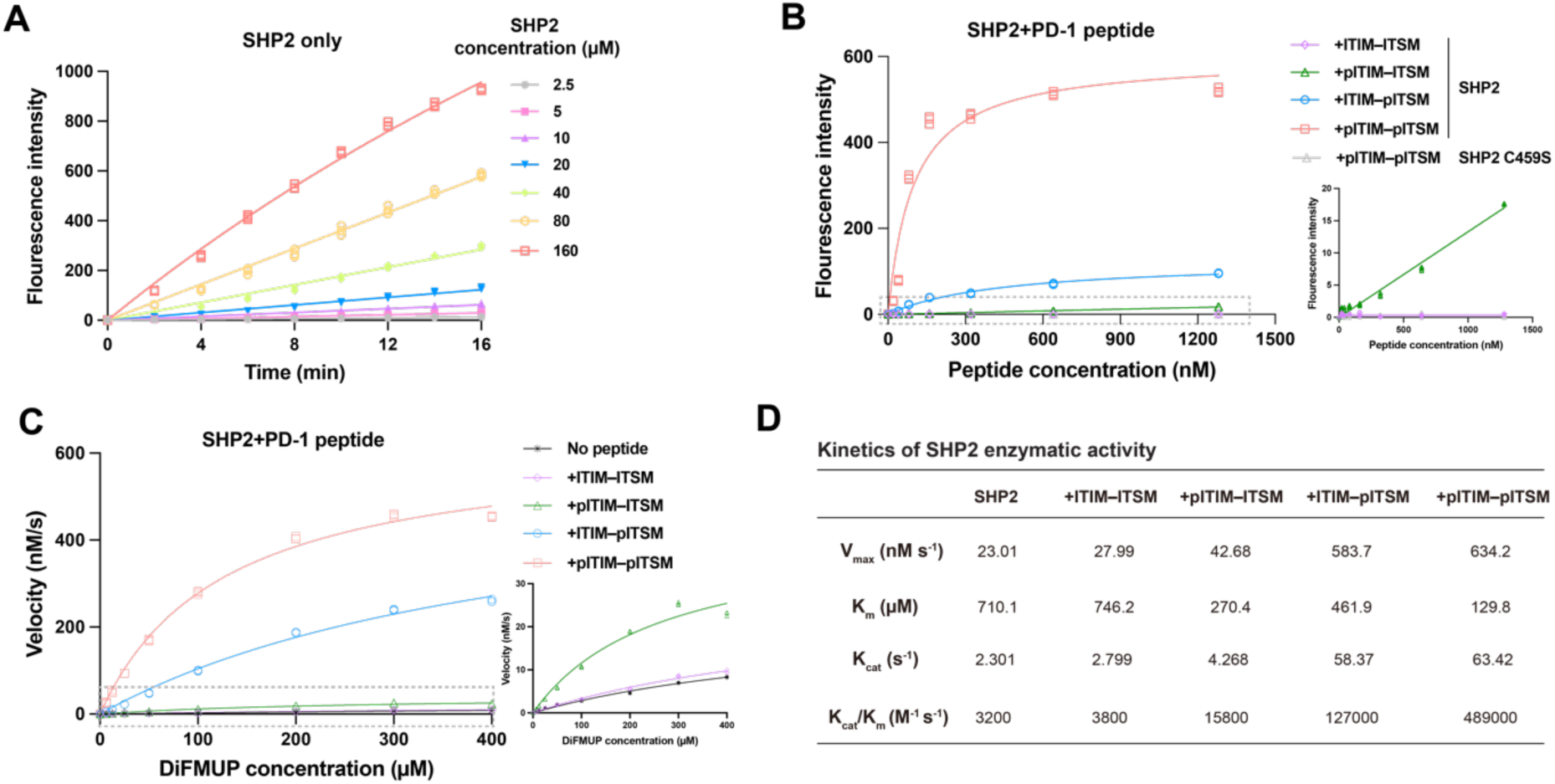
The phosphorylation of PD-1 regulates SHP2 enzymatic activity. Related to Figure 4. (A) The time- and concentration-dependent enzymatic activity of SHP2. 20 µM DiFMUP was shown (*N* = 3, *n* = 3). (B) The enzymatic activity of SHP2 in the presence of increasing amounts of phosphorylated PD-1 was measured at the 5-minute time point. 10 nM SHP2 and 20 µM DiFMUP were shown (*N* = 3, *n* = 3). (C) The PD-1 phosphorylation-mediated enzymatic velocity of SHP2 was measured at the 5-minute time point. 10 nM SHP2 and 1 µM PD-1 motif were shown (*N* = 3, *n* = 3). (D) The measured kinetics of SHP2 enzymatic activity. SHP2 shown here is core SHP2 (residues: 1–525). (p)ITIM–(p)ITSM motif (residues: 218–256, sequence: FSVD(p)YGELDFQWREKTPEPPVPCVPEQTE(p)YATIVFPS). pITSM motif (residues: 242–256, sequence: VPEQTEpYATIVFPSG). pITIM motif (residues: 218–234, sequence: VFSVDpYGELDFQWREKT).

**Figure S9.**
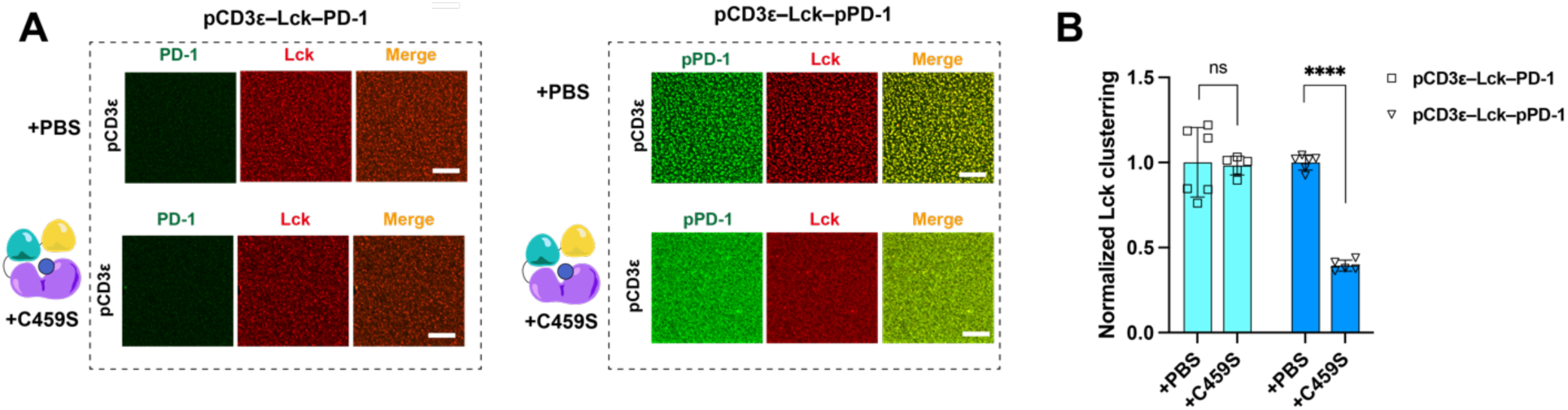
Non-catalytic SHP2 disrupts the LLPS formed by Lck–pCD3ε–pPD-1. Related to Figure 5. (A) Representative confocal images of LLPS formed by Lck–pCDε–pPD-1 with the addition of SHP2 C459S or PBS. The more phosphorylation of PD-1, the more integration of PD-1 into LLPS. (B) The quantified Lck clustering in LLPS with C459S or PBS. C459S significantly disrupts the LLPS depending on the phosphorylation of PD-1. (*N* = 3, *n* = 5 or 6). pCD3ε shown here is the intracellular region (residues: 153–207). PD-1 shown here is the intracellular region (residues: 192–288). pPD-1 was generated through the phosphorylation of PD-1 by the kinase domain of Lck. Lck shown here is the region that contains the Unique, SH3, and SH2 domains. SHP2 shown here is core SHP2 (residues: 1–525). Data in (B) were presented as mean ± SD. The significance of the difference was determined by an unpaired two-tailed Student’s *t*-test. Scale bar size: 10 µm.

**Figure S10.**
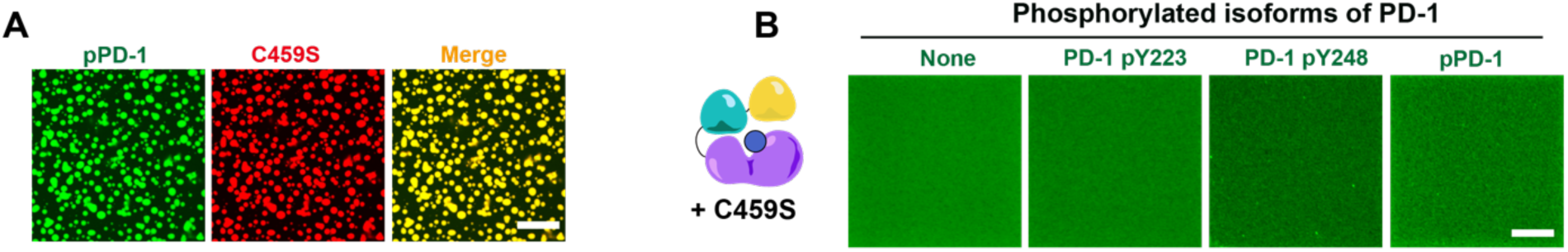
C459S binds to phosphorylated PD-1 but not to form LLPS on the SLBs. Related to Figure 5. (A) High concentration of phosphorylated PD-1 with C459S with PEG in solution. (B) Low concentration of PD-1 in different phosphorylated states with C459S on SLBs. The C459S shown here is core SHP2 (residues: 1–525) with the C459S mutation. PD-1 shown here is the PD-1 intracellular region (residues: 192–288). Scale bar size: 10 µm.

**Figure S11.**
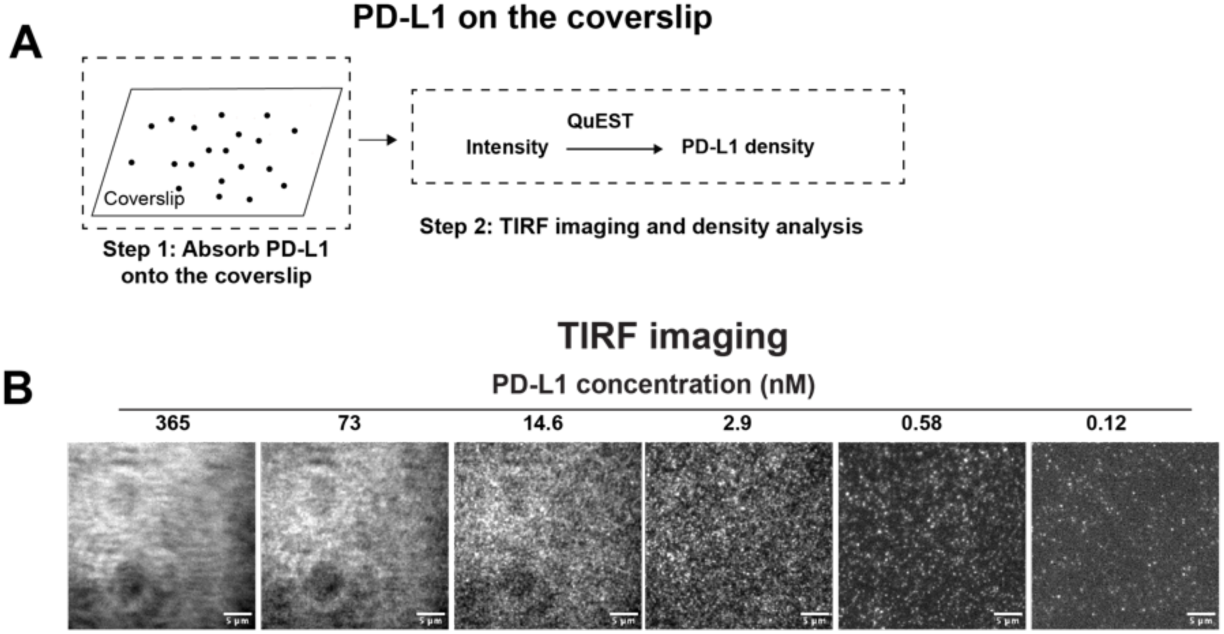
Quantification of the density of PD-L1 on the coverslip. Related to Figure 5. (A) The processes of experiment and quantification. Low concentration of PD-1 in different phosphorylated states with C459S on SLBs. (B) The representative TIRF images of PD-L1 AF405 on the coverslip.

**Figure S12.**
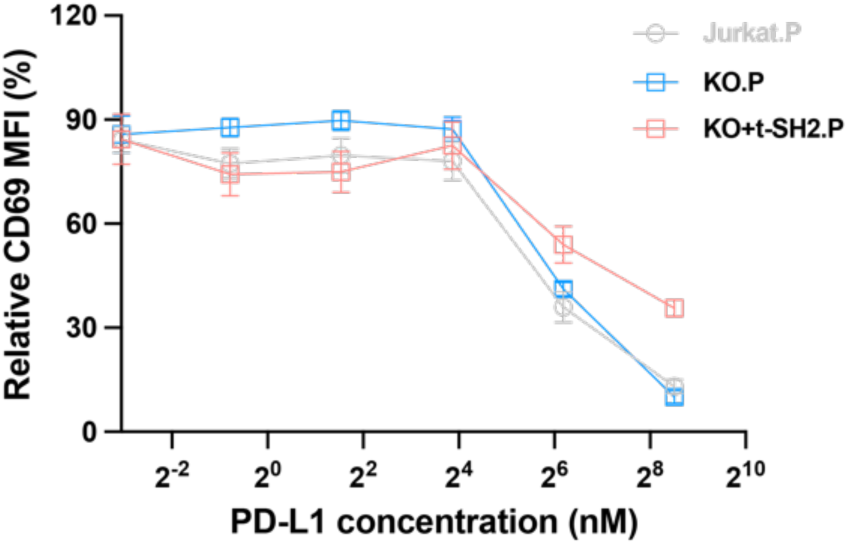
The PD-1–PD-L1 signaling-regulated activation of Jurkat, SHP2 KO, and KO+t-SH2 cells. Related to Figure 5. (*N* = 3, *n* = 6–8).

**Figure S13.**
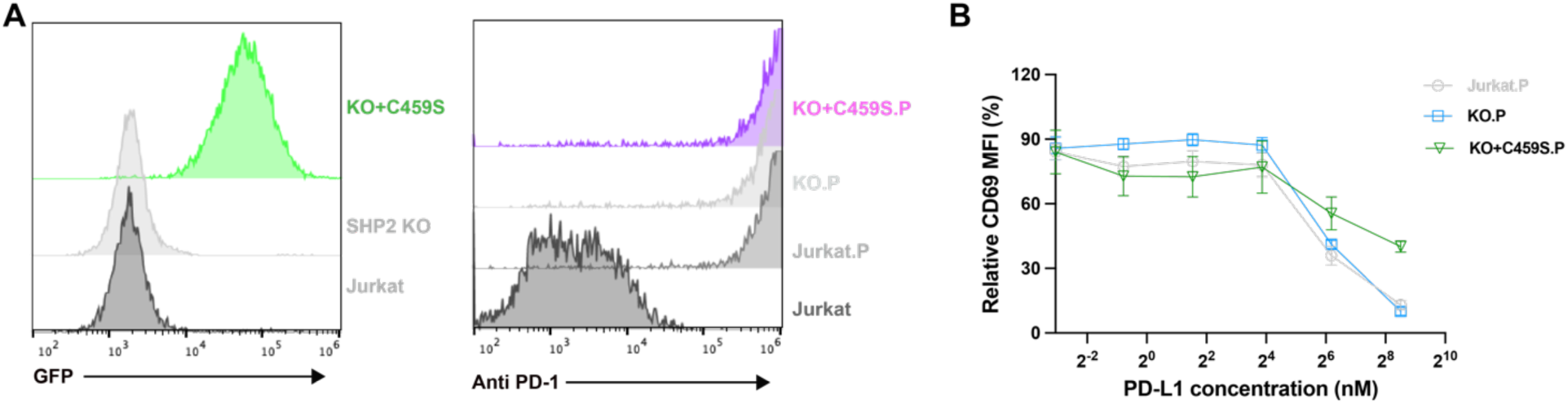
Generation of KO+C459S.P Jurkat T cells and the PD-1–PD-L1 signaling-regulated cell activation. Related to Figure 5. (A) Generation of KO+C459S.P Jurkat cells. (B) The relative CD69 MFI of engineered Jurkat T cells with different PD-L1 concentrations on the surface. (*N* = 3, *n* = 8).

## References

1. Freeman, R.M., Jr., Plutzky, J., and Neel, B.G. (1992). Identification of a human src homology 2-containing protein-tyrosine-phosphatase: a putative homolog of Drosophila corkscrew. Proc. Natl. Acad. Sci. U S A 89, 11239–11243. 10.1073/pnas.89.23.11239.

2. Adachi, M., Sekiya, M., Miyachi, T., Matsuno, K., Hinoda, Y., Imai, K., and Yachi, A. (1992). Molecular cloning of a novel protein-tyrosine phosphatase SH-PTP3 with sequence similarity to the src-homology region 2. FEBS Lett. 314, 335–339. 10.1016/0014-5793(92)81500-l.

3. Feng, G.S., Hui, C.C., and Pawson, T. (1993). SH2-containing phosphotyrosine phosphatase as a target of protein-tyrosine kinases. Science 259, 1607–1611. 10.1126/science.8096088.

4. Hof, P., Pluskey, S., Dhe-Paganon, S., Eck, M.J., and Shoelson, S.E. (1998). Crystal structure of the tyrosine phosphatase SHP-2. Cell 92, 441–450. 10.1016/s0092-8674(00)80938-1.

5. Tonks, N.K. (2013). Protein tyrosine phosphatases--from housekeeping enzymes to master regulators of signal transduction. FEBS J. 280, 346–378. 10.1111/febs.12077.

6. Lee, C.H., Kominos, D., Jacques, S., Margolis, B., Schlessinger, J., Shoelson, S.E., and Kuriyan, J. (1994). Crystal structures of peptide complexes of the amino-terminal SH2 domain of the Syp tyrosine phosphatase. Structure 2, 423–438. 10.1016/s0969-2126(00)00044-7.

7. Ishida, Y., Agata, Y., Shibahara, K., and Honjo, T. (1992). Induced expression of PD-1, a novel member of the immunoglobulin gene superfamily, upon programmed cell death. EMBO J. 11, 3887–3895. 10.1002/j.1460-2075.1992.tb05481.x.

8. Freeman, G.J., Long, A.J., Iwai, Y., Bourque, K., Chernova, T., Nishimura, H., Fitz, L.J., Malenkovich, N., Okazaki, T., Byrne, M.C., et al. (2000). Engagement of the PD-1 immunoinhibitory receptor by a novel B7 family member leads to negative regulation of lymphocyte activation. J Exp Med 192, 1027–1034. 10.1084/jem.192.7.1027.

9. Latchman, Y., Wood, C.R., Chernova, T., Chaudhary, D., Borde, M., Chernova, I., Iwai, Y., Long, A.J., Brown, J.A., Nunes, R., et al. (2001). PD-L2 is a second ligand for PD-1 and inhibits T cell activation. Nat Immunol 2, 261–268. 10.1038/85330.

10. Nishimura, H., Nose, M., Hiai, H., Minato, N., and Honjo, T. (1999). Development of lupus-like autoimmune diseases by disruption of the PD-1 gene encoding an ITIM motif-carrying immunoreceptor. Immunity 11, 141–151. 10.1016/s1074-7613(00)80089-8.

11. Dong, H., Strome, S.E., Salomao, D.R., Tamura, H., Hirano, F., Flies, D.B., Roche, P.C., Lu, J., Zhu, G., Tamada, K., et al. (2002). Tumor-associated B7-H1 promotes T-cell apoptosis: a potential mechanism of immune evasion. Nat Med 8, 793–800. 10.1038/nm730.

12. Keir, M.E., Butte, M.J., Freeman, G.J., and Sharpe, A.H. (2008). PD-1 and its ligands in tolerance and immunity. Annu Rev Immunol 26, 677–704. 10.1146/annurev.immunol.26.021607.090331.

13. Okazaki, T., and Honjo, T. (2007). PD-1 and PD-1 ligands: from discovery to clinical application. Int Immunol 19, 813–824. 10.1093/intimm/dxm057.

14. Hui, E.F., Cheung, J., Zhu, J., Su, X.L., Taylor, M.J., Wallweber, H.A., Sasmal, D.K., Huang, J., Kim, J.M., Mellman, I., and Vale, R.D. (2017). T cell costimulatory receptor CD28 is a primary target for PD-1-mediated inhibition. Science 355, 1428–1433. 10.1126/science.aaf1292.

15. Yokosuka, T., Takamatsu, M., Kobayashi-Imanishi, W., Hashimoto-Tane, A., Azuma, M., and Saito, T. (2012). Programmed cell death 1 forms negative costimulatory microclusters that directly inhibit T cell receptor signaling by recruiting phosphatase SHP2. J Exp Med 209, 1201–1217. 10.1084/jem.20112741.

16. Moore, E.K., Strazza, M., and Mor, A. (2022). Combination Approaches to Target PD-1 Signaling in Cancer. Front. Immunol. 13, 927265. 10.3389/fimmu.2022.927265.

17. Okazaki, T., Maeda, A., Nishimura, H., Kurosaki, T., and Honjo, T. (2001). PD-1 immunoreceptor inhibits B cell receptor-mediated signaling by recruiting src homology 2-domain-containing tyrosine phosphatase 2 to phosphotyrosine. Proc. Natl. Acad. Sci. U S A 98, 13866–13871. 10.1073/pnas.231486598.

18. Barford, D., and Neel, B.G. (1998). Revealing mechanisms for SH2 domain mediated regulation of the protein tyrosine phosphatase SHP-2. Structure 6, 249–254. 10.1016/s0969-2126(98)00027-6.

19. LaRochelle, J.R., Fodor, M., Vemulapalli, V., Mohseni, M., Wang, P., Stams, T., LaMarche, M.J., Chopra, R., Acker, M.G., and Blacklow, S.C. (2018). Structural reorganization of SHP2 by oncogenic mutations and implications for oncoprotein resistance to allosteric inhibition. Nat. Commun. 9, 4508. 10.1038/s41467-018-06823-9.

20. Marasco, M., Berteotti, A., Weyershaeuser, J., Thorausch, N., Sikorska, J., Krausze, J., Brandt, H.J., Kirkpatrick, J., Rios, P., Schamel, W.W., et al. (2020). Molecular mechanism of SHP2 activation by PD-1 stimulation. Sci. Adv. 6, eaay4458. 10.1126/sciadv.aay4458.

21. Jenkins, E., Fellermeyer, M., Heraghty, D.F., Sharma, S., Mitra, T., Kotowski, M., Clarke, J., Lui, Y., Daly, S., Marin, Z., et al. (2026). PD-1 signaling and PD-1 blockade-mediated tumor control are established at microvillar T cell contacts. Sci. Immunol. 11, eadz4983. 10.1126/sciimmunol.adz4983.

22. Lin, C.C., Suen, K.M., Jeffrey, P.A., Wieteska, L., Lidster, J.A., Bao, P., Curd, A.P., Stainthorp, A., Seiler, C., Koss, H., et al. (2022). Receptor tyrosine kinases regulate signal transduction through a liquid-liquid phase separated state. Mol Cell 82, 1089–1106 e1012. 10.1016/j.molcel.2022.02.005.

23. Wu, P., Zhang, T., Liu, B., Fei, P., Cui, L., Qin, R., Zhu, H., Yao, D., Martinez, R.J., Hu, W., et al. (2019). Mechano-regulation of Peptide-MHC Class I Conformations Determines TCR Antigen Recognition. Mol Cell. 10.1016/j.molcel.2018.12.018.

24. Hu, W., Zhang, Y., Fei, P., Zhang, T., Yao, D., Gao, Y., Liu, J., Chen, H., Lu, Q., Mudianto, T., et al. (2021). Mechanical activation of spike fosters SARS-CoV-2 viral infection. Cell Res 31, 1047–1060. 10.1038/s41422-021-00558-x.

25. Lei, Y., Fei, P., Song, B., Shi, W., Luo, C., Luo, D., Li, D., Chen, W., and Zheng, J. (2022). A loosened gating mechanism of RIG-I leads to autoimmune disorders. Nucleic Acids Res. 50, 5850–5863. 10.1093/nar/gkac361.

26. Chen, Y.N., LaMarche, M.J., Chan, H.M., Fekkes, P., Garcia-Fortanet, J., Acker, M.G., Antonakos, B., Chen, C.H., Chen, Z., Cooke, V.G., et al. (2016). Allosteric inhibition of SHP2 phosphatase inhibits cancers driven by receptor tyrosine kinases. Nature 535, 148–152. 10.1038/nature18621.

27. Koirala, D., Yangyuoru, P.M., and Mao, H. (2013). Mechanical affinity as a new metrics to evaluate binding events. Rev Anal Chem 32, 197–208. 10.1515/revac-2013-0004.

28. Huang, J., Zarnitsyna, V.I., Liu, B., Edwards, L.J., Jiang, N., Evavold, B.D., and Zhu, C. (2010). The kinetics of two-dimensional TCR and pMHC interactions determine T-cell responsiveness. Nature 464, 932–936. 10.1038/nature08944.

29. Patsoukis, N., Duke-Cohan, J.S., Chaudhri, A., Aksoylar, H.I., Wang, Q., Council, A., Berg, A., Freeman, G.J., and Boussiotis, V.A. (2020). Interaction of SHP-2 SH2 domains with PD-1 ITSM induces PD-1 dimerization and SHP-2 activation. Commun. Biol. 3, 128. 10.1038/s42003-020-0845-0.

30. Philips, E.A., Liu, J., Kvalvaag, A., Morch, A.M., Tocheva, A.S., Ng, C., Liang, H., Ahearn, I.M., Pan, R., Luo, C.C., et al. (2024). Transmembrane domain-driven PD-1 dimers mediate T cell inhibition. Sci. Immunol. 9, eade6256. 10.1126/sciimmunol.ade6256.

31. Fei, P., Ding, H., Duan, Y., Wang, X., Hu, W., Wu, P., Wei, M., Peng, Z., Gu, Z., and Chen, W. (2021). Utility of TPP-manufactured biophysical restrictions to probe multiscale cellular dynamics. Bio-Des. and Manuf. 4, 776–789. 10.1007/s42242-021-00163-2.

32. Dustin, M.L., Starr, T., Varma, R., and Thomas, V.K. (2007). Supported planar bilayers for study of the immunological synapse. Curr. Protoc. Immunol. Chapter 18, 18.13.11–18.13.35. 10.1002/0471142735.im1813s76.

33. Gee, K.R., Sun, W.C., Bhalgat, M.K., Upson, R.H., Klaubert, D.H., Latham, K.A., and Haugland, R.P. (1999). Fluorogenic substrates based on fluorinated umbelliferones for continuous assays of phosphatases and beta-galactosidases. Anal Biochem 273, 41–48. 10.1006/abio.1999.4202.

34. Zhu, G., Xie, J., Kong, W., Xie, J., Li, Y., Du, L., Zheng, Q., Sun, L., Guan, M., Li, H., et al. (2020). Phase Separation of Disease-Associated SHP2 Mutants Underlies MAPK Hyperactivation. Cell 183, 490–502 e418. 10.1016/j.cell.2020.09.002.

35. Kamphorst, A.O., Wieland, A., Nasti, T., Yang, S., Zhang, R., Barber, D.L., Konieczny, B.T., Daugherty, C.Z., Koenig, L., Yu, K., et al. (2017). Rescue of exhausted CD8 T cells by PD-1-targeted therapies is CD28-dependent. Science 355, 1423–1427. 10.1126/science.aaf0683.

36. Mizuno, R., Sugiura, D., Shimizu, K., Maruhashi, T., Watada, M., Okazaki, I.M., and Okazaki, T. (2019). PD-1 Primarily Targets TCR Signal in the Inhibition of Functional T Cell Activation. Front. Immunol. 10, 630. 10.3389/fimmu.2019.00630.

37. Chan, W., Cao, Y.M., Zhao, X., Schrom, E.C., Jia, D., Song, J., Sibener, L.V., Dong, S., Fernandes, R.A., Bradfield, C.J., et al. (2023). TCR ligand potency differentially impacts PD-1 inhibitory effects on diverse signaling pathways. J Exp Med 220. 10.1084/jem.20231242.

38. Chen, H., Xu, X., Hu, W., Wu, S., Xiao, J., Wu, P., Wang, X., Han, X., Zhang, Y., Zhang, Y., et al. (2023). Self-programmed dynamics of T cell receptor condensation. Proc Natl Acad Sci U S A 120, e2217301120. 10.1073/pnas.2217301120.

39. Chen, H., Wu, S., Zhang, Y., Hu, W., Lou, C., Chen, W., and Lou, J. (2026). CD28 Co-stimulation Is Organized by Signaling Condensates and Counteracted by PD-1. bioRxiv.

40. Fei, P., and Dustin, M.L. (2026). Quantitative extrapolation from single-tags (QuEST) immunofluorescence microscopy to derive TCR signalosome stoichiometries in human primary T cells. bioRxiv, 2026.2003.2028.715001. 10.64898/2026.03.28.715001.

41. Zhao, Y., Lee, C.K., Lin, C.H., Gassen, R.B., Xu, X., Huang, Z., Xiao, C., Bonorino, C., Lu, L.F., Bui, J.D., and Hui, E. (2019). PD-L1:CD80 Cis-Heterodimer Triggers the Co-stimulatory Receptor CD28 While Repressing the Inhibitory PD-1 and CTLA-4 Pathways. Immunity 51, 1059–1073 e1059. 10.1016/j.immuni.2019.11.003.

42. Xu, X., Hou, B., Fulzele, A., Masubuchi, T., Zhao, Y., Wu, Z., Hu, Y., Jiang, Y., Ma, Y., Wang, H., et al. (2020). PD-1 and BTLA regulate T cell signaling differentially and only partially through SHP1 and SHP2. J Cell Biol 219. 10.1083/jcb.201905085.

