## Supplementary Table 1 for "PD-1 remodels SHP2 dynamics and drives the non-catalytic inhibition of T cell activation"

### Table 1. Data collection and refinement statistics

| Data Set | SHP2–pPD1 complex PDB:21BJ |
|---|---|
| Space group | *P212121* |
| *a*, *b*, *c* (Å) | 65.38, 107.73, 216.68 |
| $\alpha$, $\beta$, $\gamma$ (°) | 90.0, 90.0, 90.0 |
| Resolution (Å) | 50.0 - 2.6 (2.69 - 2.60) |
| $R_{sym}$ | 0.049 (0.84) |
| $R_{pim}$ | 0.02 (0.37) |
| I/σ(I) | 35.1 (1.5) |
| $CC^{1/2}$ | 0.995 (0.81) |
| Completeness (%) | 98.0 (95.7) |
| Redundancy | 6.2 (5.8) |
| Refinement | |
| Resolution (Å) | 50 - 2.6 |
| No. of reflections | 46,483 |
| $R_{work}/R_{free}$ (%) | 23.1/26.1 |
| Protein atoms | 8,392 |
| Solvent atoms | 6 |
| B factors (Å2) | 99.3 |
| protein | 99.4 |
| Rmsd bond lengths (Å) | 0.007 |
| Rmsd bond angles (°) | 1.06 |

| | |
|---|---|
| Ramachandran outliers (%): | 0 |
| Ramachandran favored (%): | 96.1 |

Values in parentheses are for the highest-resolution shell.
