## Supplementary material for "PD-1 remodels SHP2 dynamics and drives the non-catalytic inhibition of T cell activation": STAR Methods

#### Key resources table

| REAGENT OR RESOURCE | SOURCE | IDENTIFIER |
|---|---|---|
| **Antibodies** | | |
| anti-human TCR Vβ13.1 APC | BioLegend | Cat#362407; RRID: AB_2728348 |
| anti-human CD8α PE | BioLegend | Cat#300907; RRID: AB_314111 |
| anti-human PD-1 PE | BioLegend | Cat#329905; RRID: AB_940481 |
| anti-human PD-1 Alexa Fluor 488 | BioLegend | Cat#367407; RRID: AB_2566677 |
| Rabbit monoclonal anti-human SHP2 Alexa Fluor 647 | Abcam | Cat#ab310059; Clone: Y478 |
| anti-Myc PE | Santa Cruz Biotechnology | Cat#sc-40 PE; RRID: AB_627268 |
| anti-Flag PE | Biolegend | Cat#637309; RRID: AB_2563147 |
| Rabbit monoclonal anti-human SHP2 | Santa Cruz Biotechnology | Cat#sc-280; RRID: 632401 |
| Goat anti-Rabbit IgG IRDye 800CW | LI-COR | Cat#926-32211; RRID: AB_621843 |
| anti-human PD-1 Alexa Fluor 647 | BioLegend | Cat#367420; RRID: AB_2721354 |
| anti-human CD69 Alexa Fluor 647 | BioLegend | Cat#310906; RRID: AB_314841 |
| Rabbit monoclonal anti-human phosphorylated CD28 (Tyr191) | Cell Signaling Technology | Cat#16399T; Clone: E5B9Z |
| F(ab)-Goat anti-rabbit IgG Fc Alexa Fluor 647 | Thermo Fisher Scientific | Cat#A66788 |
| anti-human CD3ε | BioXCell | Cat#BE0231; RRID: AB_2687713; Clone: UCHT-1 |
| **Bacterial and virus strains** | | |
| DH5 alpha chemocompetent cells | Vazyme | Cat#C502-02 |
| BL21 (DE3) chemocompetent cells | Vazyme | Cat#C504-02 |
| Lentivirus expressing human 1G4 TCRαβ | This study | N/A |
| Lentivirus expressing human CD8αβ | This study | N/A |
| Lentivirus expressing human PD-1 WT | This study | N/A |
| Lentivirus expressing human PD-1 Y223F | This study | N/A |
| Lentivirus expressing human PD-1 Y248F | This study | N/A |
| Lentivirus expressing human PD-1 Y223F Y248F | This study | N/A |
| Lentivirus expressing human SHP2 t-SH2–GFP | This study | N/A |
| Lentivirus expressing human SHP2 C459S–GFP | This study | N/A |
| **Chemicals, motifs, and recombinant proteins** | | |
| Chemical: SHP099 | MedChemExpress | Cat#HY-100388 |
| Chemical: Dimethylsulfoxide (DMSO) | Thermo Fisher Scientific | Cat#20688 |

| | | |
|---|---|---|
| Chemical: 6,8-difluoro-4-methylumbelliferyl phosphate (DiFMUP) | Thermo Fisher Scientific | Cat#D6567 |
| Chemical: (3-Aminopropyl)triethoxysilane (APTES) | Sigma–Aldrich | Cat#440140 |
| Chemical: Glutaraldehyde solution | Sigma–Aldrich | Cat#354400 |
| Chemical: Alexa Fluor 405 dye | Thermo Fisher Scientific | Cat#A30000 |
| Chemical: DOPC lipid | Avanti Polar Lipids | Cat#850375 |
| Chemical: Biotin-PE lipids | Avanti Polar Lipids | Cat#860511 |
| Chemical: Ni–NTA lipid | Avanti Polar Lipids | Cat#790404 |
| Chemical: biotin–PEG3500–NHS | JenKem Technology | Cat#A5026 |
| Chemical: RPMI 1640 medium | Gibco, | Cat#31870074 |
| Chemical: DMEM medium | ThermoFisher | Cat#11965092 |
| Chemical: Opti-MEM medium | Gibco | Cat#11058021 |
| Chemical: Streptavidin-coated magnetic bead | Thermo Fisher Scientific | Cat#65305 |
| Chemical: polystyrene bead | Polyscience | Cat#17145-5 |
| Chemical: Epoxy-labelled beads powder | Thermo Fisher Scientific | Cat#14301 |
| Chemical: Quantum R-PE MESF bead | Bangs Labs | Cat#827A |
| Chemical: CRISPR–Cas9 enhancer | Integrated DNA Technology | Cat#1075916 |
| Peptide: pITIM–pITSM | QYAOBIO | N/A |
| Peptide: pITIM–ITSM | QYAOBIO | N/A |
| Peptide: ITIM–pITSM | QYAOBIO | N/A |
| Peptide: ITIM–ITSM | QYAOBIO | N/A |
| Peptide: pITIM | QYAOBIO | N/A |
| Peptide: pITSM | QYAOBIO | N/A |
| Peptide: pCD3ε | GL Biochem | N/A |
| Peptide: pCD28 | GL Biochem | N/A |
| Protein: Avi–10×His–SHP2–Flag–Spy | This study, for MT experiment | N/A |
| Protein: Avi–10×His–SHP2 E76A–Flag–Spy | This study, for MT experiment | N/A |
| Protein: Avi–10×His–N-SH2 C-SH2–Flag–Spy | This study, for MT experiment | N/A |
| Protein: Avi–10×His–PTP–Flag–Spy | This study, for MT experiment | N/A |
| Protein: 10×His–spy-catcher | This study, for MT experiment | N/A |
| Protein: Avi–Myc–PD-1 | This study, for micropipette adhesion experiment | N/A |
| Protein: Avi–Myc–PD-1 Y223F | This study, for micropipette adhesion experiment | N/A |
| Protein: Avi–Myc–PD-1 Y248F | This study, for micropipette adhesion experiment | N/A |
| Protein: Avi–Myc–PD-1 Y223F Y248F | This study, for micropipette adhesion experiment | N/A |
| Protein: SHP2–Flag–Spy | This study, for micropipette adhesion experiment | N/A |
| Protein: t-SH2–Flag–Spy | This study, for micropipette adhesion experiment | N/A |

| | | |
|---|---|---|
| Protein: Lck | This study, for in vitro PD-1 phosphorylation experiment | N/A |
| Protein: Lck kinase domain | This study, for in vitro PD-1 phosphorylation experiment | N/A |
| Protein: 10×His–PD-1 | This study, for LLPS experiment | N/A |
| Protein: 10×His–PD-1 Y223F | This study, for LLPS experiment | N/A |
| Protein: 10×His–PD-1 Y248F | This study, for LLPS experiment | N/A |
| Protein: 10×His–PD-1 Y223F Y248F | This study, for LLPS experiment | N/A |
| Protein: SHP2 | This study, for enzymatic activity, and LLPS experiments | N/A |
| Protein: SHP2 C459S | This study, for crystal structure, ITC, MST, enzymatic activity, and LLPS experiments | N/A |
| Protein: N-SH2 | This study, for ITC and MST experiments | N/A |
| Protein: C-SH2 | This study, for ITC and MST experiments | N/A |
| Protein: t-SH2 | This study, for ITC, MST, and LLPS experiments | N/A |
| Protein: 10×His–UD–SH3–SH2 | This study, for LLPS experiment | N/A |
| Protein: 9C–HLA–A2–10×His–Avi | Wu et al., 2019, for SLB experiment | https://www.cell.com/molecular-cell/fulltext/S1097-2765(18)31072-4 |
| Protein: ICAM-1 ECD–Avi–12×His | This study, for SLB experiment | N/A |
| Protein: CD86 ECD–Avi–12×His | This study, for SLB experiment | N/A |
| Protein: PD-L1 ECD–Avi–12×His | This study, for SLB experiment | N/A |
| Protein: UCHT-1 Fab–12×His AF568 | Fei et al., 2026, for SLB experiment | https://www.biorxiv.org/content/10.64898/2026.03.28.715001v1 |
| Protein: PD-L1 ECD–12×His AF405 | Fei et al., 2026, for SLB experiment | https://www.biorxiv.org/content/10.64898/2026.03.28.715001v1 |
| Protein: Cas9 | Thermo Fisher Scientific | Cat#A36496 |
| **Critical commercial assays** | | |
| ClonExpress II One Step Cloning Kit | Vazyme | Cat#C113 |
| Gel DNA Extraction Mini Kit | Axygen | Cat#AP-MN-P-250 |
| Plasmid Extraction Mini Kit | Axygen | Cat#AP-GX-250 |
| JCSG Core I–IV | QIAGEN | Cat#130920 |
| Crystal Screen | Hampton Research | Cat#HR2-110 |
| Index Crystallization Screen | Hampton Research | Cat#HR2-144 |
| Quantum™ MESF beads | Bangs Labs | Cat#827A |
| **Deposited data** | | |

| | | |
|---|---|---|
| Table S1, Crystal structure of SHP2 complexed with doubly phosphorylated PD-1 motifs | This study | PDB: 21BJ |
| **Experimental models: Cell lines** | | |
| HEK293T cells | American Type Culture Collection (ATCC) | Cat#CRL-3216 |
| Jurkat E6-1 cells | American Type Culture Collection (ATCC) | Cat#TIB-152 |
| Jurkat 76 cells | Kindly provided by Dr. Jie Sun's lab at Zhejiang University | N/A |
| SF9 cells | National Biomedical Cell Resource (BMCR) | Cat#1101INS-PUMC000117 |
| 1G4 TCR, CD8, PD-1-expressing J76 cells | This study | N/A |
| 1G4 TCR, CD8, PD-1 Y223F-expressing J76 cells | This study | N/A |
| 1G4 TCR, CD8, PD-1 Y248F-expressing J76 cells | This study | N/A |
| 1G4 TCR, CD8, PD-1 Y223F Y248F-expressing J76 cells | This study | N/A |
| SHP2 KO Jurkat cells | This study | N/A |
| SHP2 KO, SHP2 t-SH2-expressing Jurkat cells | This study | N/A |
| SHP2 KO, SHP2 C459S-expressing Jurkat cells | This study | N/A |
| **Oligonucleotides** | | |
| 5' GCGCACTGGTGATGACAAAG 3' | Integrated DNA Technology | SHP2 KO CRISPR–Cas9 sgRNA |
| 5' CATTGGGATCACCATCGTGT 3' | Integrated DNA Technology | SHP2 KO CRISPR–Cas9 sgRNA |
| **Recombinant DNA** | | |
| pGEX-6p-1 Avi–10×His–SHP2–Flag–Spy | This study, for MT experiment | N/A |
| pGEX-6p-1 Avi–10×His–SHP2 E76A–Flag–Spy | This study, for MT experiment | N/A |
| pGEX-6p-1 Avi–10×His–N-SH2 C-SH2–Flag–Spy | This study, for MT experiment | N/A |
| pGEX-6p-1 Avi–10×His–PTP–Flag–Spy | This study, for MT experiment | N/A |
| pGEX-6p-1 10×His–spy-catcher | This study, for MT experiment | N/A |
| pET28a CD86–10×His–Avi | This study, for TIRF experiment | N/A |
| pET28a PD-L1–10×His–Avi | This study, for TIRF experiment | N/A |
| pGEX-6p-1 GST–TEV–Avi–Myc–PD-1 | This study, for micropipette adhesion experiment | N/A |
| pGEX-6p-1 GST–TEV–Avi–Myc–PD-1 Y223F | This study, for micropipette adhesion experiment | N/A |
| pGEX-6p-1 GST–TEV–Avi–Myc–PD-1 Y248F | This study, for micropipette adhesion experiment | N/A |
| pGEX-6p-1 GST–TEV–Avi–Myc–PD-1 Y223F Y248F | This study, for micropipette adhesion experiment | N/A |
| pGEX-6p-1 SHP2–GST–TEV–Flag–Spy | This study, for micropipette adhesion experiment | N/A |

| | | |
|---|---|---|
| pGEX-6p-1 N-SH2 C-SH2–GST–TEV–Flag–Spy | This study, for micropipette adhesion experiment | N/A |
| pFastBac 1 6×His–MBP–3C–Lck | This study, for in vitro PD-1 phosphorylation experiment | N/A |
| pFastBac 1 6×His–MBP–3C–Lck kinase domain | This study, for in vitro PD-1 phosphorylation experiment | N/A |
| pET28a 10×His–PD-1 | This study, for LLPS experiment | N/A |
| pET28a 10×His–PD-1 Y223F | This study, for LLPS experiment | N/A |
| pET28a 10×His–PD-1 Y248F | This study, for LLPS experiment | N/A |
| pET28a 10×His–PD-1 Y223F Y248F | This study, for LLPS experiment | N/A |
| KS 6×His–SUMO–C-SH2 | This study, for ITC and MST experiments | N/A |
| KS 6×His–SUMO–N-SH2 C-SH2 | This study, for ITC, MST, and LLPS experiments | N/A |
| pET28a 10×His–Lck UD–SH3–SH2 | This study, for LLPS experiment | N/A |
| pMD2.G | Addgene | Cat#12259 |
| psPAX2 | Addgene | Cat#12260 |
| pHAGE CD8αβ | This study, for lentivirus infection | N/A |
| pHAGE 1G4 TCRαβ | This study, for lentivirus infection | N/A |
| pHAGE PD-1 | This study, for lentivirus infection | N/A |
| pHAGE PD-1 Y223F | This study, for lentivirus infection | N/A |
| pHAGE PD-1 Y248R | This study, for lentivirus infection | N/A |
| pHAGE PD-1 Y223F Y248F | This study, for lentivirus infection | N/A |
| **Software and algorithms** | | |
| ImageJ | Schneider et al.[1] | https://fiji.sc |
| Prism 10 | GraphPad Software | https://www.graphpad.com |
| Labview 2020 | National Instruments (NI) | https://www.ni.com/labview |
| Adobe Illustrator 2025 | Adobe Inc. | https://www.adobe.com/products/illustrator.html |
| PyMOL | The PyMOL Molecular Graphics System | https://www.pymol.org |
| Chimera | UCSF | https://www.cgl.ucsf.edu/chimera/ |
| NanoTemper Analysis | NanoTemper Technologies | https://nanotempertech.com/ |
| XDS | Kabsch | https://xds.mr.mpg.de/ |

| | | |
|---|---|---|
| Pointless | CCP4 | https://www.ccp4.ac.uk/html/pointless.html |
| Aimless | CCP4 | https://www.ccp4.ac.uk/html/aimless.html |
| Phaser | CCP4 | https://phaser.io/ |
| Coot | Emsley et al., 2004 | https://www2.mrc-lmb.cam.ac.uk/personal/pemsley/coot/ |
| PHENIX | Adams et al., 2010 | https://www.phenix-online.org/ |
| Hidden Markov Model analysis | Kindly provided by Dr. Jie Yan's Lab at University of Singapore | N/A |
| QI algorithm (modified) | van Loenhout et al., 2012 | https://www.sciencedirect.com/science/article/pii/S0006349512004535 |
| Origin 7.0 | OriginLab Corporation | https://www.originlab.com/index.aspx?go=PRODUCTS/Origin |

### Cell Lines

HEK293T and Jurkat E6-1 cell lines were purchased from the American Type Culture Collection (ATCC). Jurkat 76 cell line was kindly provided by Dr. Jie Sun's lab at Zhejiang University. SF9 cells were purchased from the National Biomedical Cell Resource (BMCR), Chinese Academy of Medical Sciences. HEK293T cells were cultured in the DMEM culture medium (DMEM medium supplemented with 10% fetal bovine serum and 1% Penicillin-Streptomycin). Jurkat or J76-derived cells E6-1 were cultured in the RPMI 1640 culture medium (RPMI 1640 medium supplemented with 10% fetal bovine serum and 1% Penicillin-Streptomycin). SF9 cells were cultured in Grace's insect medium (GIM without bovine serum albumin, fetal calf serum, and whole egg ultrafiltrate, and with 10% fetal bovine serum).

### Method details

#### Peptides synthesis

For crystal structure, MT, ITC, MST, TIRF, and enzymatic activity experiments: PD-1 peptides were synthesized by QYAOBIO. (p)ITIM motif (residues: 218–234, sequence: VFSVD(p)YGELDFQ WREKT), (p)ITSM motif (residues: 242–256, sequence: VPEQTE(p)YATIVFPSG), (p)ITIM–(p)ITSM motif (residues: 218–256, sequence: FSVD(p)YGELDFQ WREKTPEPPVPCVPEQTE(p)YATIVFPS).

For liquid–liquid phase separation experiment: pCD3ε peptide (residues: 153–207, sequence: KNRKAKAKPVTRGAGA GGRQRGQNKERPPPVPNPDpYEPIRKGQRDLpYSGLNQRRI) was synthesized by GL Biochem. pCD28 peptide (residues: 180–220, sequence: RSKRSRLLHSDpYMNMTPRRPGPTRKHYQPpYAPPRDFAAYRS) was synthesized by GL Biochem.

#### Plasmid construction, protein expression, and purification

For magnetic tweezers experiment: Human cDNA encoding SHP2 (residues: 1–525), SHP2 E76A (residues: 1–525, Glu76 was mutated to Ala76), t-SH2 (residues: 1–220), and PTP (residues: 237–529) were separately cloned and inserted into a pGEX-6p-1 vector with ClonExpress II One Step Cloning kit with N-terminal Avi–10×His tags and C-terminal flag–spy tags. cDNA encoding spy-catch was cloned following the above method but with an N-terminal 10×His tag. To minimize potential interactions between protein and the added tags, GGGSG linkers were employed to separate SHP2 from the tag or between two tags. All constructs were verified by sequencing at TSINGKE. The verified constructs were then transformed into BL21(DE3) E. Coli. and shaken at 37˚C, 220 RPM. 1 mM IPTG was added at $OD_{600}$ 0.4–0.6 for inducing protein expression, and then shaken at 16˚C, 180 RPM for 16 h. E. Coli. were harvested and then lysed by ultrasonication, the expressed protein supernatant was collected by NTA-Ni Agarose at 4°C. Protein solution was then eluted with 250 mM imidazole buffer and then diluted into 20 mM $NaH_2PO_4$, 5% glycerin, and loaded on NTA-Ni Agarose HiTrap Q HP column, being eluted with a gradient of 8-column volumes from 0–500 mM NaCl buffer. The Q HP column-purified protein was then biotinylated in buffer containing 1×PBS, 5 mM $MgCl_2$, 2 mM ATP, 1 μM BirA, and 0.15 mM D-Biotin at 30˚C for 30 min. Proteins were finally purified with gel filtration on AKTA pure and verified by SDS-PAGE.

For liquid–liquid phase separation experiment: Human cDNA encoding Lck (UD–SH3–SH2, residues: 3–226), PD-1 (intracellular region, residues: 192–288), PD-1 Y223F (intracellular region, residues: 192–288, Tyr223 was mutated to Phe223), PD-1 Y248F (intracellular region, residues: 192–288, Tyr248 was mutated to Phe248), and PD-1 Y223F Y248F (intracellular region, residues: 192–288, Tyr223 and Try248 were mutated to Phe223 and Phe248 respectively) were separately cloned and inserted into a pET28a vector with an N-terminal 10×His tag. Lck (UD–SH3–SH2) and PD-1 proteins were expressed and purified using the above method.

For the TIRF imaging experiment: Human cDNA encoding CD86 (extracellular region, residues: 1–247), PD-L1 (extracellular region, residues: 1–238), and ICAM-1 (extracellular region, residues: 1–480) were separately cloned and inserted into a pET28a vector with C-terminal Avi–12×His tags. CD86, PD-L1, and ICAM-1 were expressed, biotinylated, and purified using the above method. Human 9C-HLA-A2 protein was from our previous publication.[2]

For micropipette adhesion experiment: Human cDNA encoding PD-1 (intracellular region, residues: 192–288), PD-1 Y223F (intracellular region, residues: 192–288, Tyr223 was mutated to Phe223), PD-1 Y248F (intracellular region, residues: 192–288, Tyr248 was mutated to Phe248), and PD-1 Y223F Y248F (intracellular region, residues: 192–288, Tyr223 and Try248 were mutated to Phe223 and Phe248 respectively) were separately cloned and inserted into a pGEX-6p-1 vector with N-terminal GST–TEV–Avi–Myc tags. Human cDNA encoding SHP2 (residues: 1–525) and t-SH2 (residues 1–220) were separately cloned and inserted into a pET28a vector with N-terminal GST–TEV and C-terminal Flag–Spy tags. The expression of protein follows the above method. The expressed protein supernatant was collected by GST beads (pre-washed with $ddH_2O$ and PBS) and incubated for 2 hours by gentle rotation in GST beads binding buffer (PBS) at 4°C. To remove the GST tag, the beads were washed with TEV cleavage buffer (20 mM HEPES, pH 8.0, 150 mM NaCl, 0.1% NP-40, 1 mM EDTA) and incubated with TEV protease in the same buffer supplemented with fresh 1 mM DTT overnight at 4°C. Proteins were finally purified with gel filtration on AKTA pure and verified by SDS-PAGE. Proteins were biotinylated following the above method.

For in vitro PD-1 phosphorylation: Human cDNA encoding Lck (residues: 3–509) and the Lck kinase domain (residues: 230–509) were separately cloned and inserted into the pFastBac 1 vector with an N-terminal fusion tag consisting of a 6×His sequence, maltose-binding protein (MBP), and a 3C protease cleavage site. Recombinant baculoviruses were generated using the Bac-to-Bac expression system. After three rounds of amplification in SF9 insect cells, the amplified virus was used to infect High Five™ insect cells cultured in ESF921 medium for protein expression. Cells were harvested 48 h post-infection and resuspended in lysis buffer (20 mM Tris–HCl, 250 mM NaCl, 0.1 mM PMSF, 10% glycerol, 0.3 mM TCEP, pH 8.5). Following centrifugation, the supernatant was applied to a 5 mL MBP Sepharose column. Bound proteins were eluted with 10 mM maltose. The MBP tag was cleaved by overnight digestion with 3C protease at 4 °C. The cleavage mixture was then passed through a 3 mL Ni Sepharose column to remove the liberated

MBP and uncleaved fusion protein. Final purification was achieved by gel filtration on a Superdex$^{TM}$ 200 Increase column using an ÄKTA Pure™ system. Fractions corresponding to the target protein peak were collected and analyzed by SDS-PAGE. The purified protein was buffer-exchanged into storage buffer containing 20 mM Tris–HCl, 200 mM NaCl, and 0.3 mM TCEP, pH 8.0.

For crystal structure, ITC, MST, enzymatic activity, and liquid–liquid phase separation experiments: Human cDNA of SHP2 (residues: 1–525), SHP2 C459S (residues: 1–525), N-SH2 (residues: 4–103), C-SH2 (residues: 106–218), t-SH2 (residues: 1–218) were separately cloned and inserted into a KS vector with a 6×His and SUMO tags on the N-terminal site. The recombinant fusion protein was expressed in the E. coli strain Rosetta. The transformed cells were grown and induced by 0.4 mM IPTG. Then, harvest cells were resuspended in lysis buffer containing 20 mM Tris–HCl, pH 8.5, 250 mM NaCl, 0.1 mM PMSF, 10% glycerol, and 0.3 mM TCEP. Centrifuged cell lysate and flowed supernatant through a Ni sepharose column, the His–SUMO tag was cleaved by ULP1 and subsequently removed by a second step Ni column purification. The recombinant protein was further purified by an ion-exchange column and a Superdex 200 increase column on AKTA Pure. Protein was collected and stored in a buffer containing 20 mM Tris–HCl, pH 8.0, 150 mM NaCl, and 0.3 mM TCEP.

**Gene knockout (KO) with CRISPR–Cas9 and validation with Western Blot (WB)**

7.5 µL of 21 µM Cas9 protein was mixed with 2 µL of 100 µM grRNA (a combination of the two grRNAs listed in the Resources Table) targeting the human *PTPN11* gene, followed by incubation at 37°C for 15 min. Subsequently, 1 µL of 200 µM enhancer was added to the CRISPR–Cas9 mixture. A total of 1.5 million Jurkat cells were harvested, washed twice with Opti-MEM medium, and resuspended in Opti-MEM containing the CRISPR–Cas9 mixture to a final volume of 100 µL. The cell suspension was transferred to a 2-mm electroporation cuvette and electroporated using an ECM 830 electroporator (BTX) under the following conditions: 275 V, 2 ms, with either one pulse (KO-1 in Figure 5E) or two pulses (KO-2 in Figure 5E). Following electroporation, cells were cultured in RPMI 1640 medium at 37°C in a humidified incubator with 5% $CO_2$. After 5 days, SHP2 expression was assessed by western blotting using a rabbit monoclonal anti-human SHP2 primary antibody and a goat anti-rabbit IgG IRDye 800CW secondary antibody.

**Generation of lentivirus and cell lines**

2 µg plasmids containing the cDNA for the target protein (1G4 TCRαβ, CD8αβ, PD-1 isoforms, t-SH2–GFP, or C459S–GFP), 2 µg lentivirus packaging plasmid, psPAX2, and 2 µg lentivirus envelope plasmids, pMD2.G, were mixed with 9 µg polyethylenimine (PEI) in the 200 µL DMEM medium at room temperature for 20 min. The mixtures were then incubated with 70%-density HEK293T cells in a 6-well plate at 37°C with 5% $CO_2$ for 48 hours. Subsequently, the lentivirus-containing supernatant was harvested and concentrated. The concentrated lentivirus was then incubated with cells for 24 hours at 37°C with 5% $CO_2$. After that, the lentivirus was removed from the cells. Following an additional 48 hours of culture, 1G4 TCR, CD8, and PD-1 on cells were validated by antibody staining, t-SH2–GFP or C459S–GFP in cells were validated by GFP signal through flow cytometry. If the positive frequency is below 90%, fluorescence-activated cell sorting (FACS) was used to purify cells.

**Protein crystallization**

The freshly purified SHP2 C459S protein was mixed with pITIM and pITSM peptides in a 1:2:2 molar ratio for crystallization screening. Screen kits were purchased from QIAGEN and Hampton Research, including JCSC core I, JCSG core II, GCSG core III, GCSG core IV, Crystal Screen, and Index Crystallization Screen. SHP2 and PD-1 mixture

(0.6 µL) and reservoir solution (0.6 µL) were mixed in a 96-well plate by hanging drop vapor diffusion method. Crystals were grown in a crystallization buffer containing 70 mM Sodium acetate/Hydrochloric acid, pH 4.6, 30% (v/v) Glycerol, 5.6% (w/v) PEG 4000 at 4°C. The crystals were harvested and cryoprotected with reservoir solution supplemented with 30% glycerol and flash-frozen in liquid nitrogen for X-ray diffraction. Diffraction data were collected at beamline BL19U1 of Shanghai Synchrotron Radiation Facility in China (SSRF). Data were indexed, integrated, and scaled using the XDS, CCP4 program Pointless, and Aimless. The structure of the SHP2–pPD-1 complex was determined by molecular replacement using the SHP2 structure (PDB:2SHP) as a searching model with Phaser. The structural model was built using Coot and refined using PHENIX. The statistics of the data collection and refinement are shown in Table S1. Structural graphs were generated using PyMOL (The PyMOL Molecular Graphics System, Version 2.0, Schrödinger, LLC) and UCSF Chimera.

**Preparation of supported lipid bilayers (SLBs) and TIRF imaging of cells**

The protocol to prepare the SLBs has been published in our study.[3,4]

In Figures 4G–4I: lipid bilayers containing 96% DOPC and 4% Biotin-PE lipids in the molar ratio were used. To prepare the experimental chamber, a glass-bottom 384-well plate was immersed in a 2.5% Hellmaxin solution at 50°C for 3 hours, followed by two rinses with deionized water. Subsequently, the plate underwent two immersions in the 5 M KOH at 50°C for 15 minutes each, then was thoroughly rinsed with deionized water and equilibrated with PBS buffer. The freshly prepared lipid bilayer solution was centrifuged at 20,000 g for 1 hour, and the resulting supernatant was carefully applied to the glass-bottom well at 37°C for 1 hour, forming the single-layered SLBs, with any excess lipids being washed away. SLBs' mobility was verified by fluorescence recovery after photobleaching (FRAP). The SLBs were then blocked with the blocking buffer (PBS with 0.1% BSA) at room temperature (RT) for 30 minutes. Subsequently, 5 µg/mL of streptavidin (SA) was conjugated to the SLBs at RT for 15 minutes. Next, 2 µg/mL of biotinylated NY-ESO-1–HLA-A2, 0.5 µg/mL CD86, 0.5 µg/mL PD-L1, and 0.2 µg/mL ICAM-1 were coated onto the SLBs by incubating for 60 minutes at RT. Finally, 1G4 TCR, CD8, and PD-1-expressing J76 cells were gently deposited onto the protein-functionalized SLBs at 37°C for 30 minutes in the reaction buffer (20 mM Hepes, 137 mM NaCl, 5 mM KCl, 1 mM $CaCl_2$, 2 mM $MgCl_2$, 0.7 mM $Na_2HPO_4$, 6 mM D-Glucose, and 0.1% BSA). After that, cells were immediately fixed with 4% PFA at RT for 10 minutes, and then the cell membrane was mildly permeabilized using 0.1% saponin at RT for 10 minutes. Before staining, cells were blocked with 3% BSA at RT for 30 minutes. Subsequently, cells were incubated with Alexa Fluor 647-labelled SHP2 and Alexa Fluor 488-labelled PD-1 antibody in the reaction buffer containing 0.02% saponin at RT for 1 hour.

In Figures 5F–5J: lipid bilayers containing 87.5% DOPC and 12.5% NTA–Ni lipids in the molar ratio were used. 0.02 µg/mL UCHT-1 AF568, 0.1 µg/mL ICAM-1, 0.19 µg/mL CD86, and 0.35 or 0 µg/mL PD-L1 were coated onto SLBs at RT for 1 hour. After SLBs were blocked, cells were incubated with protein-functionalized SLBs at 37°C for 30 minutes, and then fixed and permeabilized, followed by the pCD28 antibody staining at 4°C overnight. On the second day, primary pCD28 antibodies were washed away, and cells were then incubated with F(ab)-Goat anti-rabbit IgG Fc AF647 at RT for 1 hour.

Any excess antibody was thoroughly washed away before the TIRF imaging. The TIRF imaging parameters were kept consistent across all experiments.

**In vitro PD-1 phosphorylation and validation**

25 μM PD-1 protein (intracellular domain) isoforms were incubated with 5 μM full-length Lck or the Lck kinase domain in the PBS solution containing 1 mM ATP and 10 mM $Mg^{2+}$. The incubation was carried out on ice for 2 hours. Subsequently, the protein solution was desalted, and the phosphorylation levels of PD-1 were confirmed using mass spectrometry. The increase of 80 in molecular weight was viewed as a single phosphorylation event on the tyrosine.

**Steered molecular dynamics (SMD) simulation**

The crystal structure of the human SHP2 (PDB code: 6CMP[5]) was used as an initial model for MD simulations. The missing residues of SHP2 on this crystal structure were built using the Modloop server.[6] The protein was solvated in a rectangular box of TIP3P water molecules. $Na^+$ and $Cl^-$ ions were added to neutralize the whole system (~0.15 M). The resulting systems were first pre-equilibrated to relax the added missing region of protein, counter ions, and water molecules. Subsequently, the production simulations lasted ~200 ns with a 2-fs time step. Two independent repeating simulations were performed.

Constant-velocity steered MD simulations were performed to investigate the unfolding process of SHP2, in which representative snapshots of the production run were extracted as its initial conformations. The C-terminal Cα atom of the SHP2 was constrained at its initial position with a spring of spring constant ~1400 pN/nm, and its N-terminal Cα atom was pulled with a dummy spring of spring constant ~70 pN/nm, which moved at a speed of ~0.1 nm/ns.

The systems were processed using AmberTools and the Amber ff14SB force field for proteins.[7,8] The Energy minimizations and MD simulations in this study were performed with NAMD under periodic boundary conditions.[9] During these simulations, the temperature of the systems was maintained at 310 K with Langevin dynamics, and the pressure was controlled at 1 atm with the Nosé-Hoover Langevin piston method. Particle Mesh Ewald summation was used for electrostatic calculation, and a 12 Å cutoff was used for short-range non-bounded interactions.

**Development of Magnetic tweezers (MT) and force calibration in MT**

For details about MT development and force calibration, please refer to our recently published paper.[10-12] Briefly, MT was developed based on an inverted Nikon microscope. A pair of manipulator-controlled magnets was used to apply mechanical force onto magnetic beads, which were linked to single molecules. As a consequence, the mechanical force was transferred to the single molecules without decline. The conformational changes and protein dynamics for single molecules under different conditions (e.g., force, drug, ligand, etc.) were captured with a home-made program written in LabVIEW, which was derived from the QI algorithm,[13] and the recorded data were analyzed with another home-made program, which was written in LabVIEW. To accurately apply the mechanical force at the pico-newton (pN) level onto single molecules, the force was carefully calibrated with λ-DNA according to bead thermal fluctuation.[14,15] The correlation of magnet height (*H*) and mechanical force (*F*) yielded a double exponential curve described by the following equation:

$$F=353.3*\exp^{(-3.6*H)} + 88.4*\exp^{(-0.72*H)}$$

Where *H* is the magnet's height (mm), *F* is the mechanical force (pN) applied to the magnetic bead. According to the above equation, the mechanical force was accurately output by controlling the magnet's height with an electrical manipulator.

**Single-molecule experiment with MT**

A piece of coverslip was cleaned and functionalized with an amine on the surface by immersion in 1% aminopropyltriethoxysilane (APTES) for 1 hour. It was then packed with the other piece of non-functional coverslip in a sandwich pattern to form the experimental chamber[10]. The chamber was incubated with 0.5% glutaraldehyde PBS solution at RT for 40 minutes. After that, 2 μM spy-catcher protein was incubated in the chamber at RT for 10 min, then 20 μl amino-coated polystyrene bead was flowed in for a 4-hour incubation at RT. Subsequently, the chamber was incubated with 2% BSA solution at RT for 2 h to block the potential nonspecific interaction. Next, SHP2 proteins were diluted to pM level and incubated in the chamber at RT. 20 minutes later, the solution containing magnetic beads was slightly injected into the chamber, incubating for 20 min to form independent single-molecule tethers. All single-molecule MT experiments were done at RT in the PBS buffer condition containing 1% BSA. For experiments involving SHP099 or pITIM–pITSM, 1 μM SHP099 or 1 nM pITIM–pITSM was added at the proper time points and kept with single SHP2 molecules afterwards. Upon the increasing force, the dynamic bead heights, which represent the conformations of a single molecule, were real-time tracked. The sudden bead increments were recorded as the conformational changes of the single molecule. The force vs bead height curves were presented after averaging the raw data every 20 data points in Figure 2D. Under the constant force, the dynamic bead heights were real-time tracked. The sudden bead increments were recorded as the conformational changes of the single molecule. The time vs bead height curves were presented as raw data in Figures 3B, 3C, and S5B. Moreover, the raw data were fitted with a 2-state or 3-state pattern by the Hidden Markov Chain algorithm (kindly provided by a gift from Dr. Jie Yan's lab at the National University of Singapore) to obtain the unbinding (or unfolding) kinetics ($k_u$), rebinding (or refolding) kinetics ($k_r$), and conformational change size[16]. The time vs bead height curves were presented after averaging the raw data every 50 data points in Figures 3F and S4B.

The free energy of an unbinding (or unfolding) event in a single SHP2 molecule was approximately estimated as the following equation.[17,18]

$$\Delta G^0 = F_{eq} * \Delta x - \ln(k_u/k_r)$$

Where $\Delta G^0$ is the free energy, $F_{eq}$ is the equilibrium force, and $\Delta x$ is the conformational change size. Hence, force-dependent free energy was estimated as the following equation:

$$\Delta G^F = \Delta G^0 - F * \Delta x$$

Where $\Delta G^F$ is force-dependent free energy, $F$ is force. Based on the above two equations, the SHP2 energy landscapes under different conditions (*e.g.*, force, drug, ligand, etc.) were plotted accordingly.

**Affinity of SHP2 and PD-1 measured by the micropipette adhesion assay**

The human red blood cells (RBCs) were prepared from the volunteers' fingerstick blood with the approval of the Ethics Committee of Zhejiang University. Then, RBCs were washed twice with Hepes buffer (10 mM Hepes, 145 mM NaCl, pH 7.4), and cells were biotinylated by being coated with biotin–PEG3500–NHS with rotation at RT for 30 min. After being washed with Hepes buffer containing 1% BSA, RBCs were coated with streptavidin (SA) with rotation at RT for 30 min. Extra SA was washed away, and biotinylated phosphorylated PD-1 proteins were coated to RBCs by biotin-streptavidin specific interaction by incubation at RT for 30 minutes with rotation. Epoxy-labelled bead powder was resolved into PBS buffer (pH 7.4), and a suitable spy-catcher protein was mixed with the bead solution with rotation for 24 hours at RT. The extra spy-catcher protein was washed away, and the beads were blocked with PBS buffer containing 1% BSA for 2 h. Recombinant SHP2 or t-SH2 protein with an N-terminal spy tag was coated to beads through spy tag–spy catcher covalent linkage with rotation at RT for 30 minutes. As previously described,[19,20] affinity between PD-1 and SHP2 was measured by micropipette adhesion assay. Briefly, a SHP2 or t-SH2 protein-coated bead was controlled to contact with a PD-1-coated RBC for a duration ($t$=0.1, 0.2, 0.5, 1, and 2 s), then the bead was withdrawn from the RBC, and the RBC's membrane deformation due to PD-1 and SHP2 or t-SH2 interaction was treated as an adhesion event. After 50 contact–withdrawal cycles, adhesion frequency ($P$) was obtained. Correlation of $P$ and $t$ yielded the following equation:

$$P=1-exp\{-m_{\mathrm{PD\text{-}1}}m_{\mathrm{SHP2}}A_cK_a\left(1-exp(k_{\mathrm{off}}t)\right)\}$$

Where $m_{\mathrm{PD\text{-}1}}$ and $m_{\mathrm{SHP2}}$ were the molecular densities of PD-1 and SHP2 protein on the RBC and bead surface, respectively, which were quantified by flow cytometry with Quantum MESF beads. The fitted parameters, $A_cK_a$ and $k_{\mathrm{off}}$, were the effective affinity and off-rate of PD-1 and SHP2 under force-free conditions, respectively.

**Affinity of SHP2 and PD-1 by the isothermal titration calorimetry binding assay (ITC)**

ITC was performed in a MicroCal ITC200 instrument (Malvern Panalytical) at 25°C. pITIM–pITSM peptide and SHP2 C459S, t-SH2, N-SH2, and C-SH2 protein samples were prepared in buffer containing 50 mM HEPES, pH 7.5, 100 mM NaCl, and 0.3 mM TCEP. The titration was carried out with 20 injections, reference power 10.0, spaced 150 s apart, and stirring speed 750 rpm. The acquired calorimetric titration data were analyzed with Origin 7.0 software.

**Affinity of SHP2 and PD-1 by the microscale thermophoresis assay (MST)**

MST was measured using the NanoTemper Monolith NT.115 instrument (NanoTemper Technologies). N-SH2 and C-SH2 protein samples were labeled with Alexa Fluor 488 and diluted to 200 nM using MST buffer (50 mM Tris-HCl, pH 8.0, 80 mM NaCl, 0.05% Tween 20, 0.3 mM TCEP). SHP2 protein was mixed with different titrations of pITIM or pITSM peptide and measured in the fused silica capillary (NanoTemper Technologies) at 25°C. The Nano Temper Analysis 2.3 software was used to fit the data for determining Kd.

**Measurement of SHP2 enzymatic activity**

SHP2 phosphatase activity was measured by monitoring the dephosphorylation of the synthetic surrogate substrate 6,8-difluoro-4-methylumbelliferyl phosphate (DiFMUP) to product 6,8-difluoro-4-methylumbelliferyl hydroxide (DiFMU) by SHP2 protein. Phosphatase reactions were performed at RT in a black 96-well flat-bottom polystyrene plate. The reaction buffer contains 60 mM HEPES, pH 7.5, 75 mM NaCl, 75 mM KCl, 1 mM EDTA, and 0.3 mM TCEP.

To determine the enzyme concentration linear with velocity, 20 μM DiFMUP was incubated with SHP2 protein varying from 2.5 to 160 nM. To determine the SHP2 protein phosphatase activity upon PD-1 peptide stimulation, 10 nM SHP2 protein was co-incubated with PD-1 peptide (ITIM–ITSM, pITIM–ITSM, ITIM–pITSM, pITIM–pITSM), varying from 20 to 1280 nM for 30 min at RT. Then 20 μM of DiFMUP substrate was added to the above system, and the DiFMUP dephosphorylation was measured by a microplate reader.

For kinetic analysis, 10 nM SHP2 protein was pre-incubated with 1 μM PD-1 peptide (ITIM–ITSM, pITIM–ITSM, ITIM–pITSM, pITIM–pITSM) for 30 min at RT, followed by the addition of DiFMUP varying from 6.25 to 400 μM for 30 min at RT. The fluorescence signal was monitored by a fluorescence microplate reader at 358 nm excitation and 455 nm emission. The total reaction volume was 100 μL per well. All reactions were repeated thrice. Raw data were processed with Graphpad Prism 10 software. Velocity data were fitted by the Michaelis-Menten equation to extrapolate the kinetic parameters.

**Liquid–liquid phase separation (LLPS) experiment**

2D phase separation: SLBs were prepared as previously described.[21] 1 μM AF647- or AF488-labelled and His-tagged Lck (UD–SH3–SH2) and 1 μM AF647- or AF488-labelled and His-tagged (p)PD-1 were ligated on SLBs, then 5 μM pCD3ε or pCD28 were added to induce clustering and condensation. Then, 2 μM SHP2 protein (t-SH2 or SHP2 C459S) was injected and incubated for 30 min. Imaging was conducted on an Olympus FV1200 equipped with a 60× oil immersion objective. For AF 647 labelling, 10% of proteins were labelled with C5-maleimide Alexa Fluor 647; For AF 488 labelling, 10% of proteins were labelled with C2-maleimide Alexa Fluor 488.

3D phase separation: 20 μM AF647-labelled SHP2 C459S and 20 μM AF488-labelled pPD-1 were mixed in the presence of 5% PEG4000 and incubated for 30 min at room temperature. Imaging was conducted on an Olympus FV1200 equipped with a 60× oil immersion objective. For AF 647 labelling, 10% of proteins were labelled with C5-maleimide Alexa Fluor 647; For AF 488 labelling, 10% of proteins were labelled with C2-maleimide Alexa Fluor 488.

Quantification and statistical analysis: For 2D phase separation, Lck clustering was calculated for quantifying the dissolution of pCD3ε–Lck–(p)PD-1 or pCD28–Lck–(p)PD-1 LLPS, which equals the SD divided by the mean intensity of Lck.

**Determination of PD-L1 density on the coverslip surface**

In our previous study, we developed the QuEST method to determine molecular density on surfaces.[22] Briefly, varying concentrations of PD-L1 AF405 (20, 4, 0.8, 0.16, 0.032, and 0.0064 μg/mL) diluted in PBS were incubated with a coverslip mounted in a flow channel at RT for 1 hour. After incubation, excess protein was washed away, and the proteins immobilized on the coverslip surface were imaged using TIRF microscopy. Fluorescence intensity was quantified, and the total number of individual protein molecules within the imaging field was calculated using the following equation:

$$N=I_M/I_s$$

where N is the number of molecules, $I_M$ is the total fluorescence intensity of PD-L1 AF405 molecules within the region of interest, and $I_S$ is the fluorescence intensity of a single PD-L1 AF405 molecule.

The molecular density on the surface was then calculated using the following equation:

$$D=N/A*82.64$$

where D is the density of PD-L1 in the region (molecules/μm$^2$), N is the number of molecules in the region, and A is the area of the region in pixels. In our imaging setup, one pixel corresponds to 0.11 μm, such that 1 μm$^2$ is equivalent to 82.64 pixels$^2$.

**Jurkat T cell activation and inhibition experiment**

A 100 μL mixture containing 0.1 μg/mL UCHT-1 IgG1 antibody, 1 μg/mL ICAM-1, 1 μg/mL CD86, and varying concentrations of PD-L1 (10, 2, 0.4, 0.08, 0.016, 0.0032, or 0 μg/mL) in PBS was added to a 96-well flat-bottom plate. The plate was incubated at 37°C for 2 hours to allow adsorption of antibodies and proteins onto the well surface. Subsequently, 60,000 engineered Jurkat T cells were added to the functionalized wells and incubated at 37°C in a humidified 5% CO2 incubator for 8 h. The stimulated engineered Jurkat T cells were then harvested, stained, and analyzed for the activation marker CD69 on the cell surface.

In Figures 5M and S12B, relative CD69 expression was calculated by normalizing the CD69 mean fluorescence intensity (MFI) of cells stimulated in the presence of PD-L1 to that of cells stimulated in the absence of PD-L1. In Figure

5N, data for the group with <1 PD-L1/μm$^2$ were obtained from cells stimulated with 0.0032 μg/mL PD-L1, whereas data for the group with ~200 PD-L1/μm$^2$ represent the average of cells stimulated with 0.08 and 0.4 μg/mL PD-L1.

#### Quantification and statistical analysis

The significance of the difference between two groups was determined in Prism 10 by an unpaired two-tailed Student's *t*-test: ****$P \leq 0.0001$, ***$P \leq 0.001$, **$P \leq 0.01$, *$P \leq 0.05$, and ns represents no significant difference.

### References


1. Schneider, C.A., Rasband, W.S., and Eliceiri, K.W. (2012). NIH Image to ImageJ: 25 years of image analysis. Nat. Methods *9*, 671–675. 10.1038/nmeth.2089.
2. Wu, P., Zhang, T., Liu, B., Fei, P., Cui, L., Qin, R., Zhu, H., Yao, D., Martinez, R.J., Hu, W., et al. (2019). Mechano-regulation of Peptide-MHC Class I Conformations Determines TCR Antigen Recognition. Mol Cell. 10.1016/j.molcel.2018.12.018.
3. Fei, P., Ding, H., Duan, Y., Wang, X., Hu, W., Wu, P., Wei, M., Peng, Z., Gu, Z., and Chen, W. (2021). Utility of TPP-manufactured biophysical restrictions to probe multiscale cellular dynamics. Bio-Des. and Manuf. *4*, 776–789. 10.1007/s42242-021-00163-2.
4. Dustin, M.L., Starr, T., Varma, R., and Thomas, V.K. (2007). Supported planar bilayers for study of the immunological synapse. Curr. Protoc. Immunol. *Chapter 18*, 18.13.11–18.13.35. 10.1002/0471142735.im1813s76.
5. Padua, R.A.P., Sun, Y., Marko, I., Pitsawong, W., Stiller, J.B., Otten, R., and Kern, D. (2018). Mechanism of activating mutations and allosteric drug inhibition of the phosphatase SHP2. Nat Commun *9*, 4507. 10.1038/s41467-018-06814-w.
6. Fiser, A., and Sali, A. (2003). ModLoop: automated modeling of loops in protein structures. Bioinformatics

. 2003 Dec 12;19(18):2500-1. *19*, 2500–2501.

7. Case, D.A., Aktulga, H.M., Belfon, K., Cerutti, D.S., Cisneros, G.A., Cruzeiro, V.W.D., Forouzesh, N., Giese, T.J., Gotz, A.W., Gohlke, H., et al. (2023). AmberTools. J Chem Inf Model *63*, 6183–6191. 10.1021/acs.jcim.3c01153.
8. Maier, J.A., Martinez, C., Kasavajhala, K., Wickstrom, L., Hauser, K.E., and Simmerling, C. (2015). ff14SB: Improving the Accuracy of Protein Side Chain and Backbone Parameters from ff99SB. J Chem Theory Comput *11*, 3696–3713. 10.1021/acs.jctc.5b00255.
9. Phillips, J.C., Braun, R., Wang, W., Gumbart, J., Tajkhorshid, E., Villa, E., Chipot, C., Skee, R.D., Kalé, L., and Schulten, K. (2005). Scalable Molecular Dynamics with NAMD. J. Comput. Chem. *26*, 1781–1802.
10. Wu, P., Zhang, T., Liu, B., Fei, P., Cui, L., Qin, R., Zhu, H., Yao, D., Martinez, R.J., Hu, W., et al. (2019). Mechano-regulation of Peptide-MHC Class I Conformations Determines TCR Antigen Recognition. Mol Cell *73*, 1015–1027 e1017. 10.1016/j.molcel.2018.12.018.
11. Hu, W., Zhang, Y., Fei, P., Zhang, T., Yao, D., Gao, Y., Liu, J., Chen, H., Lu, Q., Mudianto, T., et al. (2021). Mechanical activation of spike fosters SARS-CoV-2 viral infection. Cell Res *31*, 1047–1060. 10.1038/s41422-021-00558-x.
12. Lei, Y., Fei, P., Song, B., Shi, W., Luo, C., Luo, D., Li, D., Chen, W., and Zheng, J. (2022). A loosened gating mechanism of RIG-I leads to autoimmune disorders. Nucleic Acids Res. *50*, 5850–5863. 10.1093/nar/gkac361.

13. van Loenhout, M.T.J., Kerssemakers, J.W.J., De Vlaminck, I., and Dekker, C. (2012). Non-Bias-Limited Tracking of Spherical Particles, Enabling Nanometer Resolution at Low Magnification. Biophys J *102*, 2362–2371. 10.1016/j.bpj.2012.03.073.
14. Chen, H., Fu, H., Zhu, X., Cong, P., Nakamura, F., and Yan, J. (2011). Improved high-force magnetic tweezers for stretching and refolding of proteins and short DNA. Biophys J *100*, 517–523. 10.1016/j.bpj.2010.12.3700.
15. Strick, T.R., Allemand, J.F., Bensimon, D., Bensimon, A., and Croquette, V. (1996). The elasticity of a single supercoiled DNA molecule. Science *271*, 1835–1837. 10.1126/science.271.5257.1835.
16. Yao, M., Goult, B.T., Klapholz, B., Hu, X., Toseland, C.P., Guo, Y., Cong, P., Sheetz, M.P., and Yan, J. (2016). The mechanical response of talin. Nat Commun *7*, 11966. 10.1038/ncomms11966.
17. Bustamante, C., Chemla, Y.R., Forde, N.R., and Izhaky, D. (2004). Mechanical processes in biochemistry. Annu Rev Biochem *73*, 705–748. 10.1146/annurev.biochem.72.121801.161542.
18. Li, W., Chen, P., Yu, J., Dong, L., Liang, D., Feng, J., Yan, J., Wang, P.Y., Li, Q., Zhang, Z., et al. (2016). FACT Remodels the Tetranucleosomal Unit of Chromatin Fibers for Gene Transcription. Mol Cell *64*, 120–133. 10.1016/j.molcel.2016.08.024.
19. Hu, W., Zhang, Y., Sun, X., Zhang, T., Xu, L., Xie, H., Li, Z., Liu, W., Lou, J., and Chen, W. (2019). FcgammaRIIB-I232T polymorphic change allosterically suppresses ligand binding. Elife *8*, e46689, e46689. 10.7554/eLife.46689.
20. Huang, J., Zarnitsyna, V.I., Liu, B., Edwards, L.J., Jiang, N., Evavold, B.D., and Zhu, C. (2010). The kinetics of two-dimensional TCR and pMHC interactions determine T-cell responsiveness. Nature *464*, 932–936. 10.1038/nature08944.
21. Chen, H., Xu, X., Hu, W., Wu, S., Xiao, J., Wu, P., Wang, X., Han, X., Zhang, Y., Zhang, Y., et al. (2023). Self-programmed dynamics of T cell receptor condensation. Proc. Natl. Acad. Sci. U S A *120*, e2217301120. 10.1073/pnas.2217301120.
22. Fei, P., and Dustin, M.L. (2026). Quantitative extrapolation from single-tags (QuEST) immunofluorescence microscopy to derive TCR signalosome stoichiometries in human primary T cells. bioRxiv, 2026.2003.2028.715001. 10.64898/2026.03.28.715001.
